# Granzyme K drives a newly-intentified pathway of complement activation

**DOI:** 10.1101/2024.05.22.595315

**Authors:** Carlos A. Donado, A. Helena Jonsson, Erin Theisen, Fan Zhang, Aparna Nathan, Karishma Vijay Rupani, Dominique Jones, Accelerating Medicines Partnership RA/SLE Network, Soumya Raychaudhuri, Daniel F. Dwyer, Michael B. Brenner

**Author notes:** These authors contributed equally: Carlos A. Donado, A. Helena Jonsson.

## Abstract

Granzymes are a family of serine proteases mainly expressed by CD8^+^ T cells, natural killer cells, and innate-like lymphocytes^1,2^. Although their major role is thought to be the induction of cell death in virally infected and tumor cells, accumulating evidence suggests some granzymes can regulate inflammation by acting on extracellular substrates^2^. Recently, we found that the majority of tissue CD8^+^ T cells in rheumatoid arthritis (RA) synovium, inflammatory bowel disease and other inflamed organs express granzyme K (GZMK)^3^, a tryptase-like protease with poorly defined function. Here, we show that GZMK can activate the complement cascade by cleaving C2 and C4. The nascent C4b and C2a fragments form a C3 convertase that cleaves C3, allowing further assembly of a C5 convertase that cleaves C5. The resulting convertases trigger every major event in the complement cascade, generating the anaphylatoxins C3a and C5a, the opsonins C4b and C3b, and the membrane attack complex. In RA synovium, GZMK is enriched in areas with abundant complement activation, and fibroblasts are the major producers of complement C2, C3, and C4 that serve as targets for GZMK-mediated complement activation. Our findings describe a previously unidentified pathway of complement activation that is entirely driven by lymphocyte-derived GZMK and proceeds independently of the classical, lectin, or alternative pathways. Given the widespread abundance of *GZMK*-expressing T cells in tissues in chronic inflammatory diseases and infection, GZMK-mediated complement activation is likely to be an important contributor to tissue inflammation in multiple disease contexts.

## Introduction

The complement system was first described in the late 19^th^ century as a heat-labile component of serum that “complements” the antimicrobial activity of antibodies. Since then, it has become clear that this ancient immune-surveillance system has additional functions that regulate innate and adaptive immunity and maintain tissue homeostasis^4,5^. To exert its broad, multifaceted roles, the complement system has evolved three major mechanisms of activation– the classical, the lectin, and the alternative pathways. These three pathways elicit activation of organized proteolytic cascades that generate the same set of effector molecules: the anaphylatoxins C3a and C5a that trigger a range of pro-inflammatory and chemotactic responses, the opsonins C4b and C3b that label targets for clearance by phagocytosis and lowers the threshold for B cell activation, and the C5b-9 membrane attack complex (MAC) that lyses target cells. These pathways are typically activated by soluble pattern recognition receptors that survey the environment for exogenous and endogenous danger signals and, in response, activate distinct initiator proteases that unleash the complement cascade. For example, in the classical pathway the initiator proteases C1r and C1s are activated following recognition of antigen-antibody complexes or phospholipids exposed on dead cells^4,5^. While the complement system plays a central role in many protective immune responses and homeostasis, excessive complement activation triggers and/or sustains tissue damage in many age-related, inflammatory, and autoimmune diseases, including rheumatoid arthritis (RA)^6,7^. However, for many of these diseases, the pathways that contribute to tissue pathology by driving complement activation are not entirely known.

Recently, we found that CD8^+^ T cells expressing high levels of granzyme K (*GZMK*) are the predominant population of CD8^+^ T cells in many inflamed tissues, including RA synovium, ulcerative colitis and Crohn’s disease gut, and others^3^. These cells express low levels of the classic cytotoxic lymphocyte effectors granzyme B (*GZMB*) and perforin, have low cytotoxic potential, and produce high levels of tumor necrosis factor (TNF) and interferon gamma (IFNG). GZMK is also expressed by other lymphocytes predominantly found in tissues, including CD56^bright^ natural killer (NK) cells, ψ8 T cells, mucosal-associated invariant T (MAIT) cells, and invariant natural killer T cells (iNKT) cells^3,8^. The enrichment of *GZMK*-expressing lymphocytes in inflamed tissues indicates that they might contribute to tissue pathology, yet the function(s) of GZMK remains undefined.

The granzymes make up a family of highly homologous serine proteases contained within secretory granules of cytotoxic lymphocytes^1,2^. There are five human granzymes that differ based on the physico-chemical properties of the amino acid after which they cleave. Granzyme A (GZMA) and GZMK are the only granzymes with tryptase activity, cleaving peptide bonds immediately after the basic residues lysine and arginine. GZMB, the most extensively studied granzyme, is an aspase that cleaves after aspartate residues in a number of pro-apoptotic substrates. The discovery that GZMB induces target cell apoptosis gave rise to the notion that the primary function of granzymes is to induce cell death. However, it has become increasingly clear that granzymes have additional noncytolytic functions that promote inflammation through proteolytic processing of extracellular proteins. For example, GZMA and GZMB can cleave the extracellular matrix proteins fibronectin and type IV collagen, potentially contributing to increased migration of inflammatory cells into tissues^9–13^. While GZMA and GZMB are the most well studied granzymes, much less is known about how the other granzymes, often described as orphans, contribute to inflammation and pathology.

Here, we show that GZMK is an initiator protease that independently activates the entire complement cascade. By cleaving C4 and C2 into C4b and C2a, GZMK triggers formation of a C3 convertase that cleaves C3. Addition of nascent C3b molecules to C4bC2a allows formation of C5 convertases that cleave C5. Together, the active convertases generate the anaphylatoxins C3a and C5a, the opsonins C4b and C3b, and the C5b-9 MAC or terminal complement complex (TCC). In RA synovium, fibroblasts express the highest amounts of complement C2, C3 and C4 and secrete them in response to the T cell-derived cytokines IFNG and TNF, serving as substrates for GZMK-mediated complement activation. Further, in RA synovial tissue GZMK is enriched in areas with evidence of abundant complement activation. Given the abundance of GZMK-secreting CD8^+^ T cells and complement-producing cells in inflamed tissues, we suggest that GZMK drives a tissue-focused pathway of complement activation independent of the previously known complement pathways.

## Results

### GZMK is expressed by CD8^+^ T cells, NK cells, and innate-like T cells in blood and tissues

To determine which cells in human blood and tissues express *GZMK*, we performed flow cytometry and analyzed publicly available RNA-seq datasets. In human peripheral blood, GZMK protein was expressed by 38.4 ± 13.9% (mean ± standard deviation (SD)) of CD8^+^ T cells and a majority of mucosal-associated invariant T (MAIT) cells, V82 ψ8 T cells, and non-cytotoxic CD56^bright^ CD16^-^ NK cells (Fig. 1a,b and Extended Data Fig. 1a). A smaller fraction of CD4^+^ T cells mainly consisting of effector memory T cells (T_em_), and cytotoxic CD56^dim^ CD16^+^ NK cells also expressed GZMK (Fig. 1a,b and Extended Data Fig. 1b). Thus, many different lymphocytes in human blood express GZMK, about half of which are effector memory CD8^+^ T cells (Fig. 1c and Extended Data Fig. 1b).

**Fig. 1.**
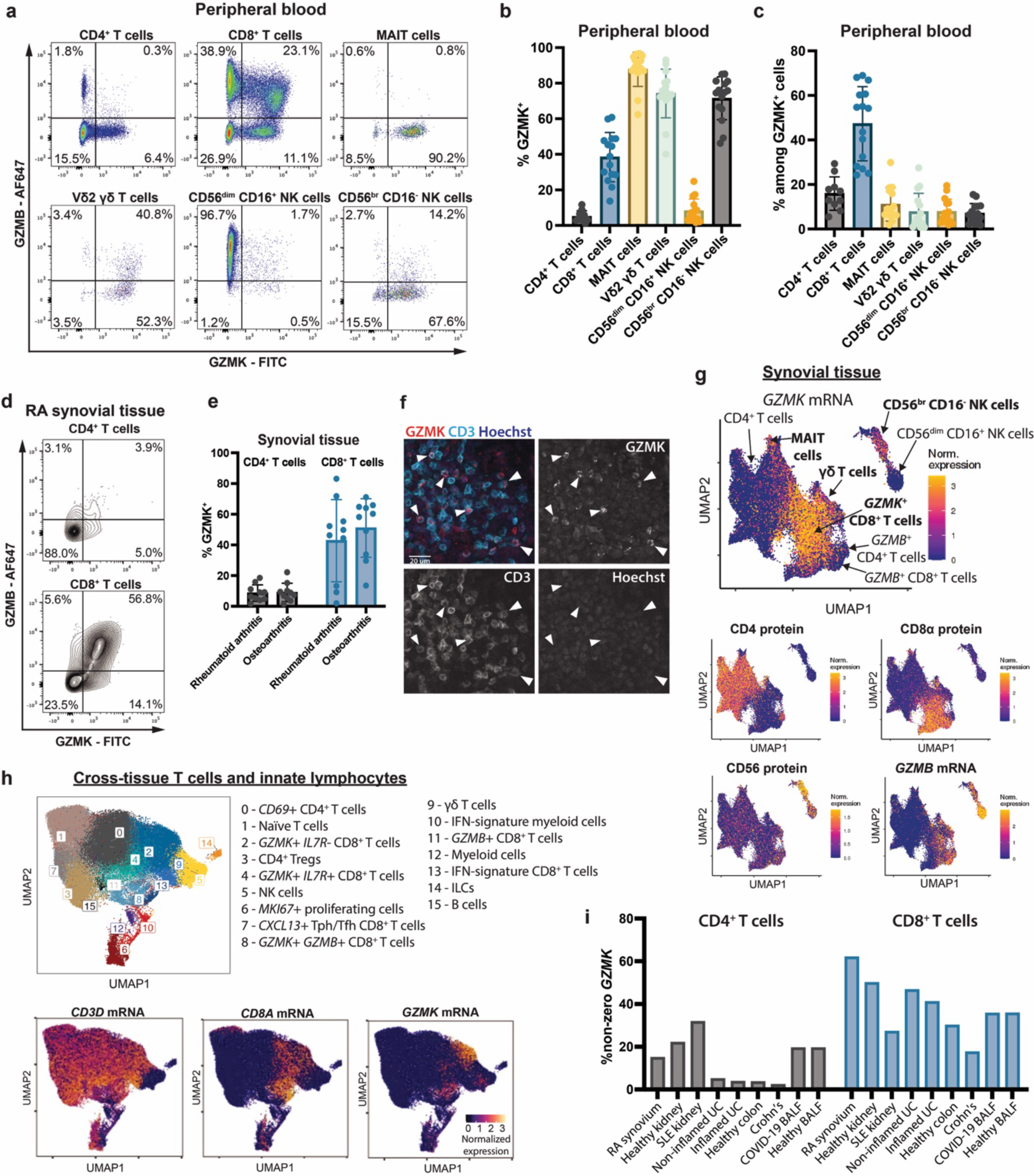
GZMK is expressed by CD8^+^ T cells, NK cells, and innate-like T cells in blood and tissues. **a,** Representative flow cytometry plots showing intracellular GZMK and GZMB staining of the indicated T cell or NK cell subset in healthy peripheral blood. **b,** Frequency of GZMK expression by the indicated T cell or NK cell subset in peripheral blood from healthy controls. **c,** Among all GZMK^+^ lymphocytes in blood, percentage belonging to each T cell or NK cell subset. In panels b and c, *n* = 10 for CD4^+^ T cells; *n* = 15 for all other populations. **d,e,** Representative flow plots and aggregate data showing expression of GZMK protein, as measured by intracellular flow cytometry of unstimulated CD4^+^ or CD8^+^ T cells from synovial tissue collected from patients with RA (*n* = 10). **f**, Representative image showing immunofluorescent staining of RA synovial tissue is shown. Arrowheads indicate examples of GZMK^+^ T cells. Data are representative of at least three independent experiments. **g,** UMAP plots displaying single-cell RNA-seq profiles from 94,056 T cells and 8,497 NK cells from synovial tissue from patients with RA (*n* = 70) or OA (*n* = 9) shaded by expression of the indicated protein marker or gene transcript. **h,** UMAP plot of Louvain clustering of 85,522 integrated single-cell RNA-seq profiles from T cells and NK cells from healthy or diseased tissues from RA synovium, Crohn’s disease (CD) ileum, ulcerative colitis colon, lupus nephritis (SLE) kidney, and COVID-19 BALF. Expression patterns of selected genes are shown for the integrative dataset in UMAP space. **i**, Percentage of cells in CD4^+^ T cell clusters (gray columns) or CD8^+^ T cell clusters (blue columns) with detectable *GZMK* gene expression, stratified by tissue and disease source. (**b,c,e)** Data are mean ± s.d.

Next, we queried GZMK expression in synovial tissue from patients with RA, a chronic autoimmune disease characterized by infiltration by T cells and other lymphocytes into affected synovial tissues. At the protein level, GZMK was expressed by half of all synovial CD8^+^ T cells in both RA and osteoarthritis (OA) (42.8 ± 26.8% vs 51.1 ± 19.1%, respectively (mean ± SD)) and about 10% of synovial CD4^+^ T cells (Fig. 1d,e and Extended Data Fig. 1c).

Immunofluorescence staining confirmed GZMK-expressing T cells were abundantly present in inflamed RA synovial tissue (Fig. 1f). For higher granularity, we assessed *GZMK* mRNA expression in a large single-cell RNA-seq dataset collected by the Accelerating Medicines Partnership: Rheumatoid Arthritis/Systemic Lupus Erythematosus (AMP RA/SLE) Network^14^. *GZMK* was expressed among a much larger number of CD8^+^ T cells than *GZMB* (Fig. 1g). In addition, we also detected *GZMK* among CD4^+^ T cells, NK cells, and ψ8 T cells (Fig. 1g and Extended Data Fig. 1d). Thus, *GZMK* is expressed by a large proportion of CD8^+^ T, NK and innate-like T cells in OA and RA synovial tissues.

To investigate the expression of *GZMK* among cells in tissues beyond the synovium, we integrated single-cell RNA-seq data from several publicly available datasets of tissue cells collected from patients with RA, lupus nephritis, ulcerative colitis, Crohn’s disease, COVID-19, and healthy controls^3,15–19^ (Fig. 1h and Extended Data Figure 2a). In these data, *GZMK* was detected in 20-60% of CD8^+^ T cells and up to 30% of CD4^+^ T cells, depending on the disease state, consistent with previously reported results^3^ (Fig. 1h,i and Extended Data Fig. 2c). In contrast, *GZMB* was detected in roughly 10-50% of CD8^+^ T cells in all tissues and fewer than 20% of CD4^+^ T cells. Collectively, our findings indicate the *GZMK*^+^ lymphocytes are highly abundant and make up the majority of CD8^+^ T in some tissues and disease settings.

### CD8^+^ T cells constitutively secrete GZMK

Next, we examined how GZMK production and release are regulated by CD8^+^ T cells. To do so, we compared the ability of peripheral blood CD8^+^ T cells to synthesize and secrete GZMK in the presence or absence of T cell receptor (TCR) stimulation. Strikingly, unstimulated CD8^+^ T cells constitutively secreted GZMK (Fig. 2a). The accumulation of GZMK in the supernatants of untreated cells was not accompanied by a concomitant decrease in intracellular levels over time, indicating that CD8^+^ T cells continuously synthesize GZMK (Fig. 2a and Extended Data Fig. 3). TCR stimulation did not lead to a significant enhancement in GZMK release. Instead, TCR crosslinking led to a decrease in intracellular GZMK (Fig. 2a and Extended Data Fig. 3), indicating TCR signaling inhibits constitutive GZMK synthesis. In contrast to GZMK, GZMB was not constitutively released in unstimulated cultured cells. Rather, it was synthesized and secreted only in response to TCR stimulation (Fig. 2a). Collectively, these data demonstrate GZMK is constitutively synthesized and released by CD8^+^ T cells in the absence of TCR stimulation. Further, this constitutive pattern of release indicates that GZMK has continuous access to extracellular substrates and is therefore able to exert functional outcomes on cells and tissues surrounding GZMK^+^ CD8^+^ T cells at all times.

**Fig. 2.**
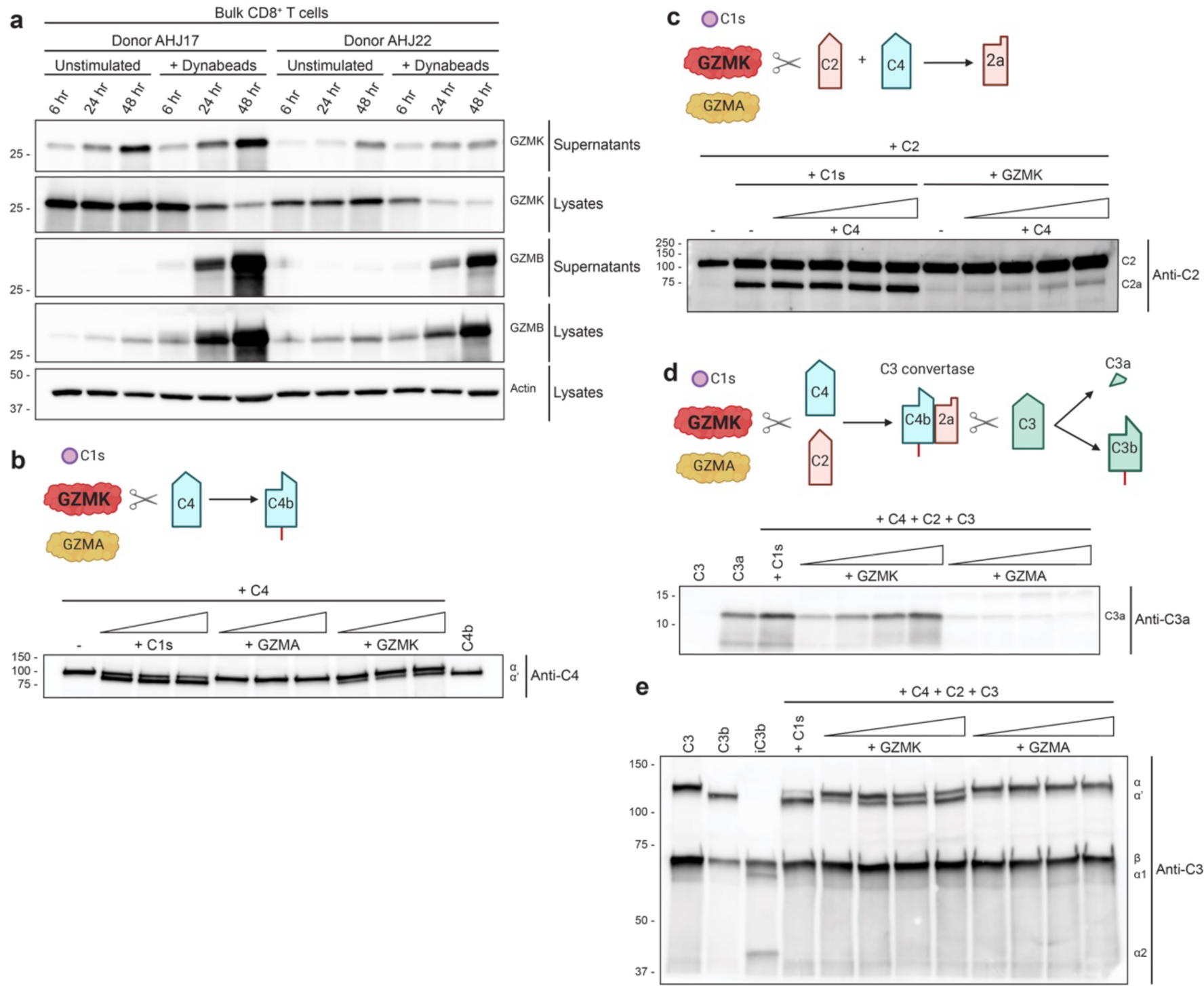
GZMK cleaves the complement components C4 and C2 to generate a C3 convertase that cleaves C3 into C3a and C3b. **a,** Bulk CD8^+^ T cells were MACS sorted from peripheral blood of two donors and either left unstimulated or stimulated with anti-CD3/CD28 dynabeads for 6, 24, or 48 hours. Precipitated supernatants and lysates were analyzed by immunoblot using antibodies against GZMK and GZMB. **b,** Increasing concentrations of active GZMK or GZMA were incubated with serum-purified C4 for 4 hours and cleavage products were analyzed by immunoblot. Active C1s was used as a positive control for C4 cleavage into C4b. Purified C4b was used to confirm the size of the C4b fragment generated by C1s and GZMK. **c,** Active GZMK was incubated with C2 for 4 hours in the presence or absence of increasing concentrations of C4 and cleavage products were analyzed by immunoblot. Active C1s was used as a positive control for C2 cleavage into C2a. **d,e,** Increasing concentrations of active GZMK or GZMA were incubated with C2 + C3 + C4 and cleavage products were analyzed by immunoblot. Active C1s was used as a positive control for generation of a C3 convertase that cleaves C3 into (**d**) C3a and (**e**) C3b. Serum-purified C3a and C3b were used to confirm the identity of the fragments generated by C1s and GZMK, while purified iC3b was used to determine whether GZMK generates inactive C3b. Schematic representation of the assays are shown for **b-e**. Data are representative of at least four independent experiments.

### GZMK cleaves complement C4 and C2 to generate active C3 convertases

The abundance of *GZMK*-expressing lymphocytes across different tissues and disease states led us to focus on understanding how GZMK might contribute to inflammation. Its constitutive release suggests that GZMK might process extracellular proteins. To identify new substrates for GZMK, we performed a protein BLAST^20^ and identified human proteins with high sequence homology to GZMK (Extended Data Fig. 4a). As expected, GZMK was most closely related to its closest homolog, GZMA. The next most homologous protein was complement factor D (CFD), the serine protease that activates the alternative complement pathway. In the alternative pathway, C3b molecules associate with complement factor B (CFB), inducing a conformational change in the latter that allows CFD to cleave CFB into a proteolytically active Bb fragment. Bb then remains complexed with C3b, forming the alternative pathway C3 convertase (C3bBb) that cleaves C3 molecules into the anaphylatoxin C3a and the opsonin C3b^5^. Based on the results of our protein BLAST, we hypothesized GZMK might also cleave CFB and, in doing so, could act as an additional mechanism to catalyze activation of the alternative complement pathway. To test this hypothesis, we used serum-purified complement components to assess whether we could recapitulate activation of the alternative pathway. While incubation of the protease CFD with CFB and C3b resulted in generation of the Bb fragment, active GZMK incubated with C3b and CFB failed to generate any Bb (Extended Data Fig. 4b).

The initiator proteases that activate the classical, lectin, and alternative complement cascades, including C1s, the mannose-binding lectin-associated serine proteases-1 and −2 (MASP-1 and MASP-2), and CFD all have one feature in common with GZMK: they are tryptase-like proteases that cleave their complement substrates after arginines or lysines^21–30^. Therefore, we next asked whether GZMK might process other proteins involved in complement activation. In contrast to the alternative pathway, the classical and lectin pathways both employ a C3 convertase composed of C4b + C2a (C4bC2a) but rely on distinct serine proteases to catalyze its generation. In the classical pathway, C1s cleaves C4 into C4b, and C2 into C2a. C4b associates with C2a to form a C3 convertase, with C2a acting as the enzymatic component that cleaves C3 into C3a and C3b^5^. Therefore, we tested if GZMK can cleave C4 and C2, using serum purified active C1s as a positive control. As expected, incubation of C4 with C1s resulted in generation of the C4b fragment (Fig. 2b). Surprisingly, GZMK independently cleaved C4 into C4b. As a specificity control, we incubated C4 with active GZMA, the only other human granzyme that can cleave peptide bonds after basic residues, but it did not generate the C4b product (Fig. 2b).

We then asked whether GZMK can cleave C2 into C2a, the active enzymatic component of the classical and lectin pathway C3 convertases. In those pathways, following cleavage of C4, C4b molecules associate with the zymogen C2 and promote its cleavage into the active form, C2a^5^. We found that incubation of C2 with GZMK in the presence of C4 resulted in cleavage of C2 into C2a (Fig. 2c), identical to the cleavage product generated by C1s. Although GZMK was not as efficient as C1s in cleaving C2, C2a is known to be a highly active protease even when present in extremely low amounts^31^. Having shown that GZMK can cleave C4 and C2 into C4b and C2a, we next asked whether those cleaved fragments assemble into a C3 convertase that can cleave C3 into C3a and C3b. To test this, we separately incubated either C1s, GZMK or GZMA with a mixture of C4, C2 and C3. Like C1s, active GZMK induced the generation of C3a (Fig. 2d) and C3b (Fig. 2e), while active GZMA did not. Addition of GZMK did not result in generation of iC3b, the inactive form of C3b (Fig. 2e). Further, GZMK did not directly cleave C3 (Extended Data Fig. 4c). These results indicate that GZMK can elicit formation of enzymatically-active C3 convertases by cleaving C4 and C2 into C4b and C2a.

### Fibroblasts are major producers of complement components that are substrates for GZMK

Multiple lines of evidence indicate that substantial production of complement components and complement activation are present in RA synovial tissue and fluid^6,32–45^. The enrichment of *GZMK*^+^ CD8^+^ T cells in tissues led us to consider local, as opposed to liver-derived complement production. To identify cell populations that could locally produce C2, C3 and C4 in RA synovium, we analyzed a publicly available bulk RNA-seq data set of T cells, B cells, monocytes, and fibroblasts from 51 samples of synovial tissue from patients with RA or OA^46^. While monocytes expressed *C2* and *C3*, fibroblasts were the largest producers of complement *C2*, *C3*, and *C4A/B* transcripts in RA synovium (Fig. 3a). Fibroblasts in RA synovial tissue also expressed C3 protein (Fig. 3b). By examining supernatants of cultured RA synovial fibroblasts, we found that resting fibroblasts secrete C2 and C3 at modest levels but not C4 (Fig. 3c,d). Given that GZMK^+^ CD8^+^ T cells are known to produce abundant IFNG and TNF^3^, we asked whether these T cell-derived cytokines regulate the secretion of C2, C3 and C4 by synovial fibroblasts. IFNG and TNF each induced C2 and C3 release (Fig. 3c,d). In contrast, only IFNG was capable of stimulating C4 release (Fig. 3c,d). Combined stimulation with IFNG and TNF resulted in a dramatic increase in C3 secretion as compared to treatment with either cytokine alone, a synergistic effect that was not observed for C2 or C4 (Fig. 3d). Thus, fibroblasts are major producers of the complement proteins C2, C3, and C4 in RA synovium, especially in response to the T cell-derived cytokines IFNG and TNF.

**Fig. 3.**
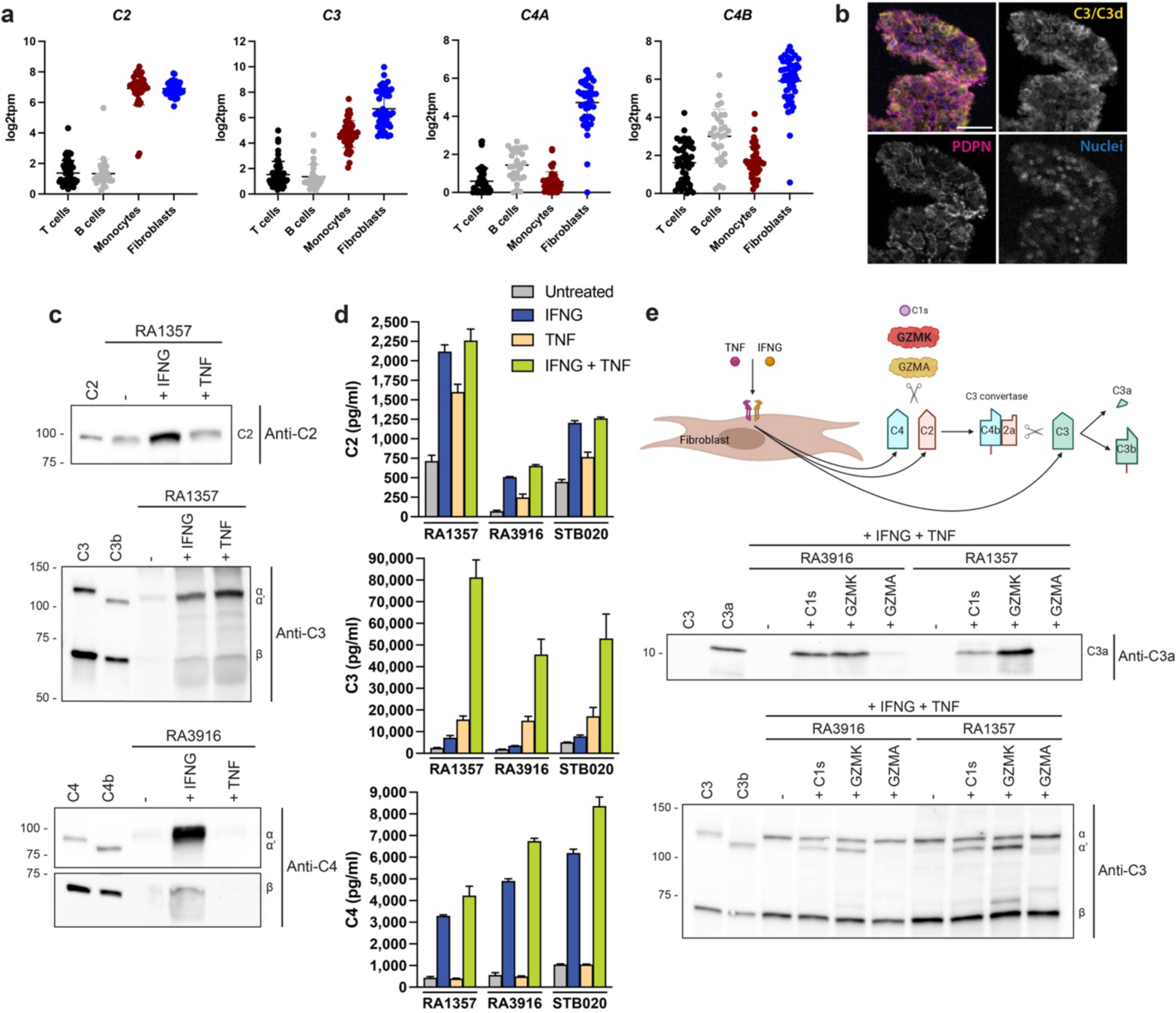
Synovial fibroblasts are major producers of tissue-derived complement proteins that are substrates for GZMK. **a,** Expression of complement *C2*, *C3*, *C4A*, and *C4B* in T cells, B cells, monocytes, and fibroblasts sorted from disaggregated synovial tissue from patients with RA (N = 33) or osteoarthritis (N = 12), measured by low-input RNA-seq using data from the AMP RA/SLE network^46^. Data are mean ± s.d. **b,** Representative image showing immunoflourescent staining of RA synovial tissue stained with antibodies against complement C3/C3d (yellow) and PDPN (magenta) and Hoechst nuclear stain (blue). Scale bar is 40 microns. **c**, Synovial fibroblasts were left untreated or stimulated with IFNG or TNF for 24 hours and the supernatants were precipitated and immunoblotted against C2, C3, or C4. Serum-purified C2, C3, C3b, C4 and C4b were run as controls to identify the proper bands. **d,** Synovial fibroblasts were left untreated or stimulated with IFNG, TNF, or IFNG + TNF for 24 hours and supernatants were assayed for the presence of C2, C3, or C4 by ELISA. Data are mean ± s.d of three technical replicates. **e,** Synovial fibroblasts were stimulated for 24 hours with IFNG + TNF after which the cell-free supernatants were left untreated or were incubated with C1s, GZMK or GZMA and cleavage products were analyzed by immunoblot for generation of C3a and C3b. Serum-purified C3a and C3b were used to confirm the identity of the fragments generated by C1s and GZMK. (**b-e**) Data are representative of at least three independent experiments. Schematic representation of the assay is shown for **e**.

Next, we tested whether active GZMK can trigger formation of an extracellular C3 convertase from complement components secreted by synovial fibroblasts. To this end, we stimulated synovial fibroblasts *in vitro* with a combination of IFNG and TNF, isolated the supernatants, and incubated them with active C1s, GZMK, or GZMA. The addition of C1s or GZMK, but not GZMA, resulted in generation of C3a and C3b (Fig. 3e) from fibroblast-derived complement components. Together, these results indicate GZMK can elicit formation of a C3 convertase by cleaving complement proteins produced by synovial fibroblasts.

### GZMK forms membrane-bound C3 convertases that generate bioactive C3a and C3b

Following processing of C3, the anaphylatoxin C3a is released into the fluid phase and acts on a number of different cell types to elicit local inflammatory responses^5^. To assess whether C3a generated through the GZMK complement activation pathway is bioactive, we tested its ability to trigger mast cell degranulation, a well-known effect of C3a^47^. To do so, we incubated active C1s or GZMK with serum purified C2 + C3 + C4 and transferred the reaction products to human mast cells in the presence of an anti-LAMP-1 antibody to detect degranulation. While the addition of C3 did not elicit mast cell degranulation, reactions containing C2 + C3 + C4 and either C1s or GZMK induced strong mast cell degranulation, comparable to that induced by serum purified C3a, as indicated by the translocation of LAMP-1 to the cell surface (Fig. 4a).

**Figure 4.**
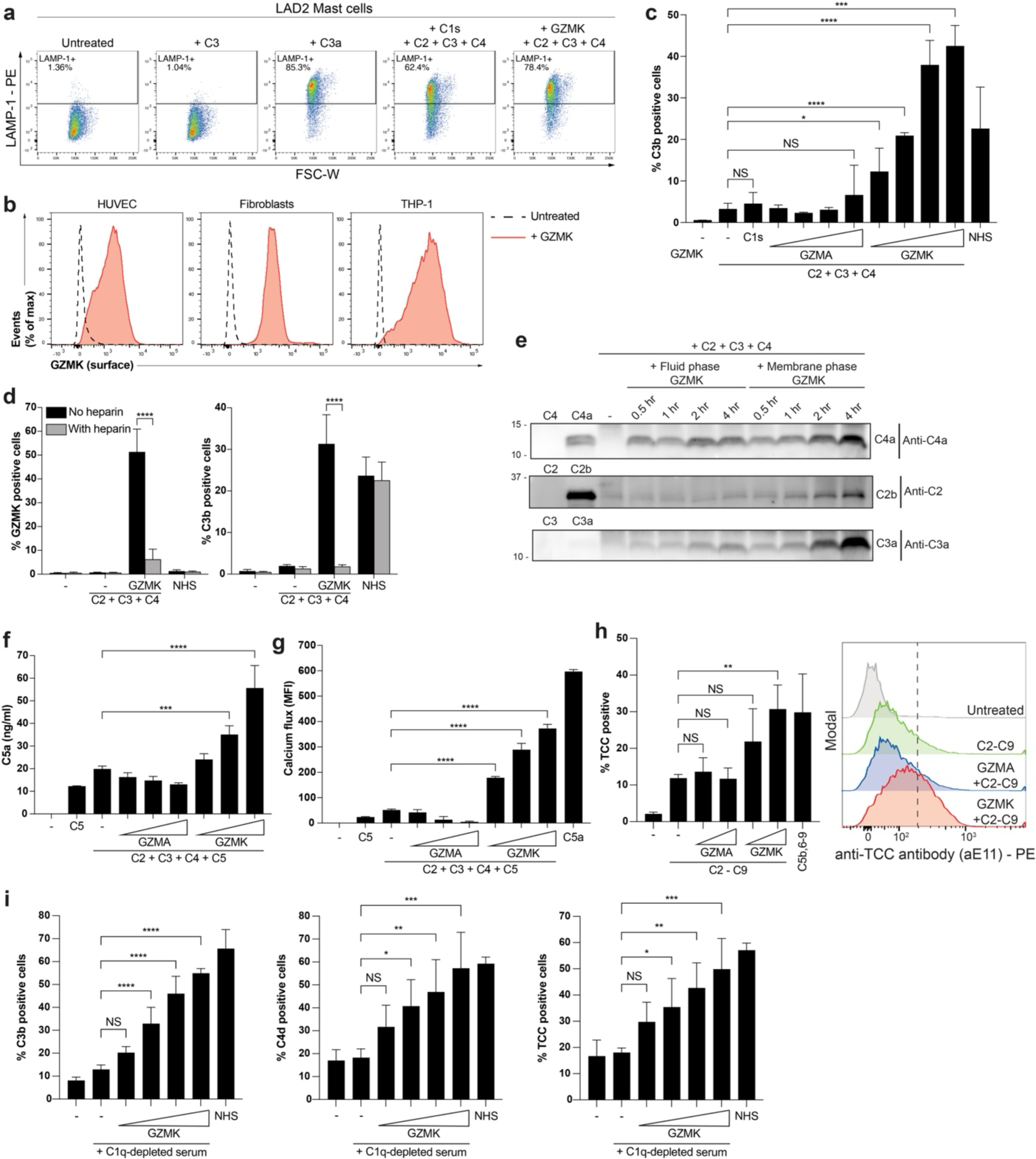
GZMK triggers activation of the entire complement cascade. **a,** LAD2 mast cells were incubated with C3, C3a, or the reaction products obtained after incubating C1s or GZMK with C2 + C3 + C4. Degranulation was then assessed by flow cytometric staining of surface LAMP-1. **b**, HUVEC cells, synovial fibroblasts, or THP-1 monocytes were incubated with recombinant GZMK for 1 hour on ice, and surface-bound GZMK was then measured by flow cytometry after staining of unfixed, unpermeabilized cells. **c**, Surface deposition of C3b on HUVEC cells after incubation with C2 + C3 + C4 with or without C1s, GZMK, or GZMA, as measured by flow cytometry. Normal human serum (NHS) was used as a positive control. **d**, GZMK surface binding (left) and C3b deposition (right) on HUVEC cells after incubation with C2 + C3 + C4 with or without GZMK in the presence or absence of heparin. NHS was used as a control. **e**, GZMK either pre-bound to the surface of HUVECs or in solution without HUVECs was incubated with C2 + C3 + C4 for 0.5, 1, 2 or 4 h and the supernatants were analyzed by immunoblot for generation of C4a, C2b, and C3a. Purified C4a, C2b, and C3a were used to confirm the size of the fragments generated by GZMK. **f,** HUVEC cells were incubated with C2 + C3 + C4 + C5 with or without GZMA or GZMK and the generation of C5a was then measured by ELISA. **g**, C5aR1-expressing Chem-1 reporter cells labeled with Fluo-5F were incubated with C5, C5a, or the supernatants obtained after incubating HUVEC cells with C2 + C3 + C4 + C5 with or without GZMA or GZMK. Calcium flux was immediately assessed by flow cytometry and is presented as MFI (MFI minus unstimulated MFI). **h**, HUVEC cells were incubated with C2 + C3 + C4 + C5 + C6 + C7 + C8 + C9 with increasing concentrations of GZMK or GZMA. C5b,6-9 was used as a positive control. Terminal complement complex formation was then measured by flow cytometry. Histograms depict representative data. **i**, Surface deposition of C3b (left) and C4d (middle), and TCC formation (right) on HUVEC cells after incubation with C1q-depleted serum and increasing concentrations of GZMK, as measured by flow cytometry. NHS was used as a positive control. Data in **a-i** are representative of at least 3 independent experiments. *P* values were calculated using (**c, f-i**) one-way analysis of variance (ANOVA) with Dunnett’s multiple comparisons tests or (**d**) a two-way ANOVA with Sidak’s multiple comparisons tests. \**P* < 0.05, \*\**P* < 0.01, \*\*\**P* < 0.001 and \*\*\*\**P* < 0.0001, NS = not significant. Data are mean ± s.d. of (**c,d,f,h**) three or (i) five independent experiments or (**g**) three technical replicates from a representative of three independent experiments. **(b)** Data are representative of three independent experiments.

Upon cleavage, newly-processed C4b and C3b fragments undergo a conformational change that exposes a highly reactive thioester that is otherwise buried in intact C4 and C3 molecules. This newly-exposed thioester mediates opsonization of targets by forming covalent bonds with nearby hydroxyl or amino groups on a surface^5^. If C4 and C3 are cleaved in the fluid phase (in solution), far from reactive groups on a surface, the thioester in C4b and C3b is rapidly inactivated by water hydrolysis, rendering it incapable of opsonizing a surface. Therefore, efficient C4b- and C3b-mediated opsonization requires that the proteases that initiate complement activation adhere to surfaces so they can cleave C4 in close proximity to reactive groups that nascent C4b molecules can covalently bind. Following cleavage of C2, the released C2a fragment can associate with this surface-bound C4b to form a membrane-bound C3 convertase that generates C3b molecules that can opsonize the same surface. To test whether GZMK can readily attach to cell membranes, we incubated human endothelial cells, human synovial fibroblasts, and THP-1 monocytes with GZMK for 1 hour on ice, washed the cells and assessed surface GZMK staining by flow cytometry. Indeed, we found abundant GZMK on the surface of nearly 100% of the cells (Fig. 4b). In contrast, serum-purified C1s was not able to bind cell membranes, except when the entire C1 complex, containing C1q, C1r, and C1s, was added to cells that were sensitized with cell type-specific antibodies that can be bound by the soluble pattern recognition receptor, C1q (Extended Data Fig. 5a). To determine if GZMK can elicit opsonization by deposition of C3b, we assessed its ability to induce covalent C3b attachment to the plasma membrane of human endothelial cells. Incubation of endothelial cells with serum-purified C2 + C3 + C4 in the presence of GZMK, but not GZMA, resulted in cell surface deposition of C3b (Fig. 4c and Extended Data Fig. 5b). In contrast, C1s was not capable of triggering C3b deposition on endothelial cells (Fig. 4c) at concentrations at which it induced generation of C3b (Fig. 2e) and bioactive C3a in the fluid phase (Fig. 2d and 4a). Therefore, in contrast to C1s, GZMK directly binds to plasma membranes to trigger formation of membrane-bound C3 convertases and to elicit efficient opsonization of target surfaces.

All human granzymes are highly cationic, with isoelectric points of about 10^1^. Further, GZMB has been demonstrated to bind plasma membranes through ionic interactions with negatively charged molecules including heparan sulfate glycosaminoglycans^48–51^. Like GZMB^51^, GZMK has a heparin binding region^52^. Thus, we tested whether GZMK associates with cell membranes by interacting with heparan sulfate glycosaminoglycans. To do so, we incubated GZMK + C2 + C3 + C4 with endothelial cells and found that addition of soluble heparin blocked the ability of GZMK to bind to cell surfaces and to induce opsonization of endothelial cells (Fig. 4d and Extended Data Fig. 5c).

Due to its strong propensity to bind membranes, we asked whether GZMK is more efficient at activating complement while it is bound to membranes or in solution. To test this, we incubated GZMK with C2 + C3 + C4 in the presence or absence of HUVECs and assessed the generation of C4a, C2b, and C3a by immunoblot. While GZMK was as efficient at cleaving C4 in the fluid and membrane phases at early time points, membrane-bound GZMK generated more C4a by the end of the reactions (Fig. 4e). Membrane-bound GZMK was also more efficient at cleaving C2 and triggered the generation of significantly more C3a over time than it did in solution (Fig. 4e). Together, these findings demonstrate that, in contrast to C1s, GZMK does not require a soluble pattern recognition receptor to direct it to a surface where it can trigger complement activation. Instead, GZMK directly binds membranes, where it is more efficient than in the fluid phase at cleaving C4 and C2, to form membrane-bound C3 convertases that generate bioactive C3a and C3b.

### GZMK elicits formation of C5 convertases to generate C5a and the TCC

Formation of a C5 convertase, the last enzymatic step in complement activation, requires addition of C3b molecules to a membrane-localized C3 convertase^53–57^. Once C3b molecules associate with C4bC2a, the substrate specificity of the convertase switches, allowing it to cleave C5 into C5a, an anaphylatoxin, and C5b^57^. The latter can associate with plasma membranes along with C6, C7, C8 and C9 molecules to form the MAC, also known as the TCC^5^. Our observations that GZMK binds cell membranes (Fig. 4b,d), triggers C3b deposition on them (Fig. 4c), and catalyzes the generation of more C4a, C2a, and C3a when bound to a membrane (Fig. 4e) suggest GZMK efficiently generates membrane-bound C3 convertases. The newly-generated C3b molecules could associate with pre-existing C3 convertases on the plasma membrane to form C5 convertases (C4bC2aC3b) that can generate C5a and the TCC. To test whether GZMK can elicit formation of a C5 convertase, we incubated endothelial cells with C2 + C3 + C4 + C5 and GZMK and assessed the generation of C5a by ELISA. GZMK, but not GZMA, triggered generation of C5a in a dose-dependent manner (Fig. 4f). The C5a detected by ELISA was bioactive, as the supernatants from endothelial cells incubated with C2 + C3 + C4 + C5 and GZMK induced calcium flux in a cell line expressing C5aR1 (Fig. 4g and Extended Data Fig. 5d). To assess whether GZMK can elicit TCC formation, we incubated endothelial cells with C2 + C3 + C4 + C5 + C6 + C7 + C8 + C9 in the presence of GZMK and evaluated the presence of membrane TCCs by flow cytometry. GZMK, but not GZMA, induced a dose-dependent increase in TCC formation on the membrane of endothelial cells comparable to the positive control, where purified C5b,6 was added to cells along with C7, C8, and C9 (Fig. 4h). GZMK-dependent TCC formation was further enhanced when endothelial cells were treated with dynasore (Extended Data Fig. 5e), an inhibitor of dynamin-dependent endocytosis, which has been reported to increase TCC formation on cell membranes^58,59^.

Our findings demonstrate that GZMK can activate the complement cascade when incubated with complement proteins purified from serum, or with the supernatants of cytokine-stimulated fibroblasts. Next, we asked whether GZMK can activate complement when incubated with whole serum. Incubation of endothelial cells with GZMK in the presence of C1q-depleted serum (devoid of antibody-mediated activation of the classical pathway) resulted in C3b and C4d (a breakdown product of C4b) deposition, as well as TCC formation, on the surface of the endothelial cells (Fig. 4i, Extended Data Fig. 5f). Together, these observations demonstrate that GZMK is capable of activating the entire complement cascade, starting with cleavage of C4 and C2, and leading to formation of C3 and C5 convertases that generate the anaphylatoxins C3a and C5a, the opsonin C3b, and the TCC.

### GZMK is enriched in regions with abundant complement activation within RA synovial tissue

To find evidence that GZMK induces complement activation in tissues, we performed immunofluorescence microscopy on RA synovial tissue sections to look for an association between areas with C3b deposition, C5a production, and GZMK. We detected complement deposition in areas where GZMK was abundant, as assessed using an antibody that detects C3d (Fig. 5 and Extended Data Fig. 6a-c), one of the breakdown products of C3b. Further, C5a was also present in regions enriched for GZMK (Fig. 5 and Extended Data Fig. 6a,d,e). We confirmed these findings using a second set of antibodies against C3d and C5a (Extended Data Fig. 6b-e). Thus, our results provide evidence consistent with abundant complement activation in GZMK-rich areas within inflamed tissues.

**Figure 5.**
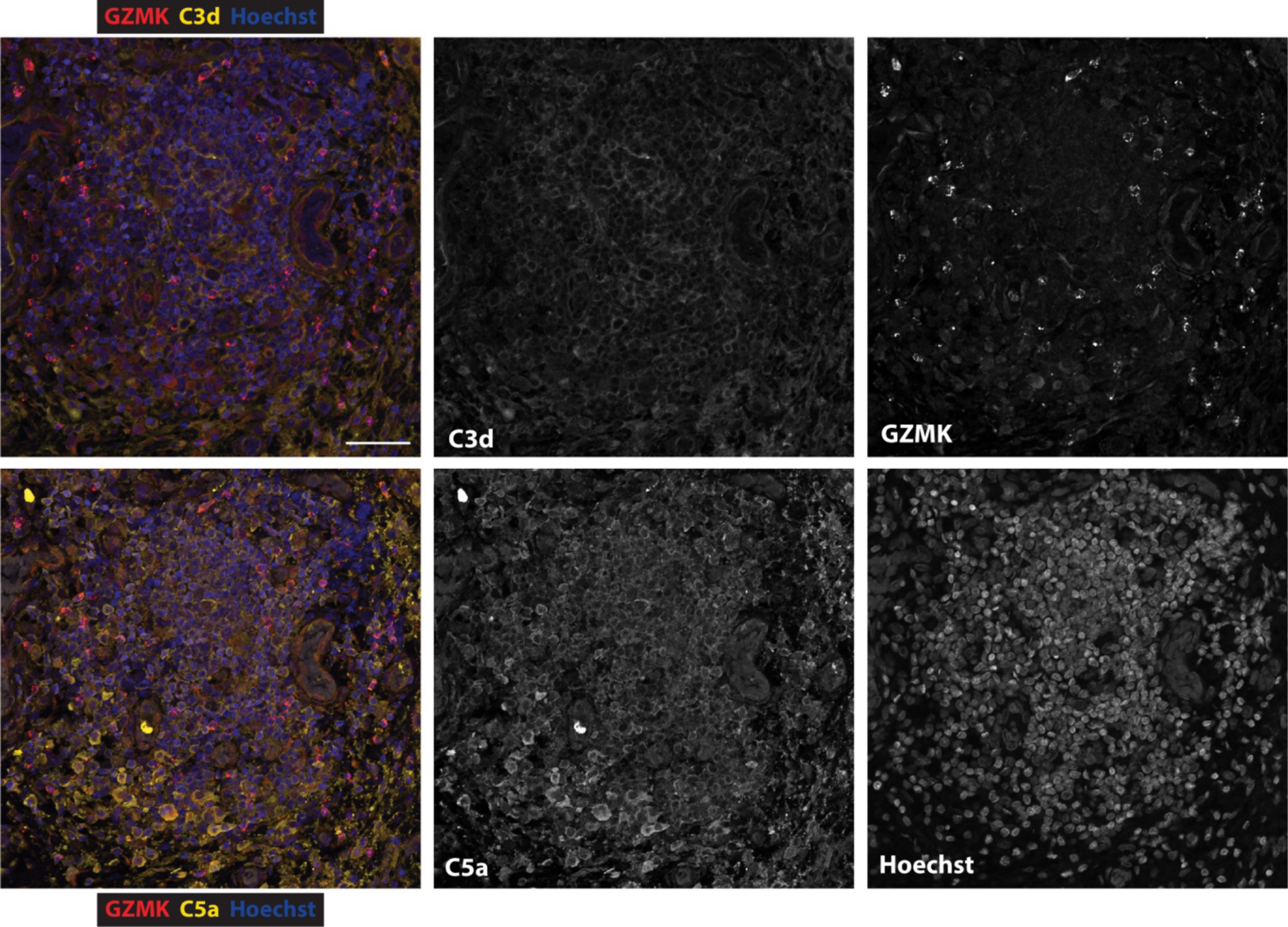
GZMK is enriched in regions with abundant complement activation within RA synovial tissue. Representative image showing immunofluorescence staining of RA synovial tissue with antibodies against C3d (clone C3D/2891, yellow in top composite), C5a (clone 2952, yellow in bottom composite), and GZMK (red) as well as Hoechst nuclear stain (blue). Scale bar represents 50 microns. Data are representative of 3 independent experiments.

## Discussion

We describe the discovery of a fourth pathway of complement activation. This previously unrecognized pathway is activated independently of the proteins that catalyze activation of the classical, lectin and alternative pathways. Instead, it is directly activated by lymphocyte-derived GZMK, a serine protease regarded as an orphan amongst granzymes due to its poorly defined function.

Akin to the tryptase-like serine proteases that catalyze activation of the classical and lectin pathways, we found that the tryptase-like GZMK protease directly cleaves C4 and C2 into C4b and C2a. While the GZMK pathway generates the same C3 and C5 convertases as the classical and lectin pathways, there are some key distinctions between them (Extended Data Fig. 7a,b). Activation of the classical and lectin pathways is initiated by the soluble pattern recognition receptors C1q and mannose-binding lectin, respectively, following their recognition of specific molecules on a target cell surface^5^. Binding to their cognate ligand on a membrane triggers two major events: 1) it activates proteolytic enzymes that are non-covalently associated to them and 2) it brings those proteases to a surface where complement activation can occur more efficiently than in solution. These proteolytic enzymes, C1s, and MASP-1 and MASP-2, act as the initiator proteases of their respective complement cascades^5^. In contrast to the initiator proteases of classical and lectin pathways, GZMK can activate the entire complement cascade independently of a soluble pattern recognition receptor. This is evidenced by several observations. First, GZMK and C1s both triggered the generation of C3a and C3b in the fluid phase following incubation with serum-purified C2 + C3 + C4 (Fig 2d,e). Second, when the same reaction was performed in the presence of endothelial cells, GZMK but not C1s induced C3b deposition on the cell membrane (Fig. 4c). Third, GZMK triggered the generation of bioactive C5a (Fig. 4f,g) and membrane-localized TCCs (Fig. 4h) when C5, C6, C7, C8 and C9 were further added to the reaction. Fourth, GZMK independently bound the membrane of several cell types (Fig. 4b,d) whereas C1s was only able to bind membranes when the entire C1 complex was added to cells that were sensitized with cell type-specific antibodies (Extended Data Fig. 5a). Together, these results demonstrate that GZMK independently binds plasma membranes, where it can orchestrate the assembly of membrane-bound C3 and C5 convertases to generate all the effector molecules of the complement cascade.

In contrast to GZMK, C1s and MASP-1/2 require the soluble pattern recognition receptors C1q and mannose-binding lectin to bind membranes and trigger full activation of the classical and lectin pathways, respectively. Due to their highly cationic nature, granzymes can associate with target cell membranes through charge-charge interactions with negatively charged glycosaminoglycans like heparan sulfate^49–51^. This receptor-independent binding mechanism could allow GZMK to direct C3 and C5 convertase formation to virtually any surface coated with heparan sulfate, whether it is a cell membrane of self or microbial origin, or onto extracellular matrix components. The lack of a need for a soluble pattern recognition receptor also allows GZMK to elicit formation of C3 convertases in the fluid phase which, although weaker than membrane-bound convertases, may serve as an important source of C3a.

Our finding that CD8^+^ T cells constitutively release GZMK suggests this complement activation pathway may be a significant contributor to inflammation wherever *GZMK*-expressing CD8^+^ T cells are abundant. Indeed, we found high percentages of *GZMK*-expressing CD8^+^ T cells across inflamed tissues, including RA synovium, Crohn’s disease ileum, and ulcerative colitis colon. While circulating complement components are largely produced by the liver, we found that in RA synovial tissue fibroblasts are the major producers of the complement substrates required for activation of this pathway. Further, IFNG and TNF, two cytokines produced abundantly by *GZMK*-expressing CD8^+^ T cells^3^, potentiate secretion of complement C2, C3, and C4 by synovial fibroblasts. It is likely that other cell types, including monocytes and macrophages, also contribute to local complement activation by providing substrates for GZMK.

In addition to the enrichment we have demonstrated in tissues affected by chronic inflammatory diseases, *GZMK*-expressing T cells have recently been found in other autoimmune diseases including Sjogren’s syndrome^60^, multiple sclerosis^61^, systemic sclerosis^62^, IgG4-related disease^63,64^, dermatomyositis^65^, and granulomateous uveitis^66^. Further, *GZMK*-expressing CD8^+^ T cells have also been reported to accumulate with age across different tissues in mice and humans^67^. These T cells may contribute to inflammaging, the increase in inflammation that occurs with aging. Indeed, *GZMK*-expressing T cells are significantly expanded in inflammaging-associated conditions, including human atherosclerotic plaques^68,69^, Alzheimer’s disease^70^, and Parkinson’s disease^71^. Further, these cells are enriched in the cancer-associated tissues of patients with colorectal cancer^72,73^, head and neck squamous cell carcinoma^74^, melanoma^75,76^, non-small cell lung cancer^77^, and others^78–88^. Given that the complement system plays an important role in atherosclerosis^89^, neurodegeneration^90^, cancer^91,92^, and other conditions linked to inflammaging, our findings raise the possibility that GZMK might contribute to inflammaging by triggering activation of the complement cascade. We also found CD8^+^ T cells expressing *GZMK* made up a significant portion of the CD8^+^ T cells in healthy tissues like kidney, colon, and lung. As previously described of the complement system, GZMK-mediated complement activation might also contribute to cellular and tissue homeostasis in healthy tissues where *GZMK*-expressing lymphocytes are present. In addition to CD8^+^ T cells, we found innate-like lymphocytes including most MAIT cells, iNKT cells, V82 ψ8 T cells and CD56^bright^ NK cells also express GZMK, and are enriched in various tissues. Therefore, GZMK-mediated complement activation may occur in many tissue contexts in health, disease, and aging. We found evidence of complement activation in regions where GZMK was enriched within RA synovial tissue, and we anticipate the same will be true across different tissues and disease states.

In summary, we have defined a pathway of complement activation mediated by GZMK that proceeds independently of soluble pattern recognition receptors or the initiator proteases of the classical, lectin, or alternative pathways. Our work highlights GZMK as a promising target to inhibit complement activation across multiple disease indications where *GZMK*-expressing lymphocytes are enriched. Unlike antibodies or inhibitors that block components common to all complement pathways, selective inhibition of GZMK has the potential to spare the anti-microbial functions of complement while specifically inhibiting this lymphocyte-enabled pathway in chronically inflamed tissues.

## Acknowledgements

This work was supported by US NIH grant no. R01 AR073290 and R01 AR081792 (M.B.B.), Rheumatology Research Foundation grant no. 889234 (M.B.B.), NIAMS K08 AR081412 (A.H.J.), Rheumatology Research Foundation Investigator Award (A.H.J.), and US NIH grant no. 5T32AR007098-48 (E.T). We thank V. Michael Holers (University of Colorado, Denver) for valuable input and discussions. S.R. is supported by NIH grants (5P01AI148102-03, 5UC2AR081023-02, 5R01AR063759-08).

## Author contributions

C.D., A.H.J., and M.B.B. conceived the study. C.D. designed, performed, and analyzed complement cleavage assays, C3a-mediated degranulation assays, C3b and C4d deposition and TCC formation assays, C5a quantification and bioactivity assays, and fibroblast assays. A.H.J. designed, performed, and analyzed GZMK expression analysis of lymphocytes, T cell stimulation flow cytometric assays, and synovial tissue microscopy studies. E.T. performed C3b deposition and TCC formation assays. C.D. and E.T. conducted GZMK surface binding studies. C.D. and K.V.R. performed and analyzed T cell stimulation flow cytometric assays. D.J. performed and analyzed synovial tissue microscopy studies. A.N. performed computational data analysis of T and NK cell subsets in RA. F.Z. performed computational data analysis of T cells across disease states. AMP provided unpublished single-cell transcriptional and proteomic data from a cohort of patients with RA and osteoarthritis. S.R. supervised the computational analyses, including the processing and analysis of unpublished data from AMP. D.D. provided cell lines and advice on experimental design and analysis. C.D., A.H.J. and M.B.B wrote the manuscript with assistance from all other authors. M.B.B. supervised experiments and data analysis.

## Additional information

Correspondence and requests for materials should be addressed to Michael B. Brenner.

## Methods

### Cell lines

Synovial fibroblast cell lines were derived from RA patients. LAD2 cells were provided by Daniel F. Dwyer. THP-1 cells were provided by the American Type Culture Collection (ATCC). HUVEC cells were provided by Lonza. All cells were cultured in a humified, 5% CO_2_ incubator at 37 °C. Fibroblasts were grown in DMEM supplemented with 2 mM L-glutamine, 100 U/ml penicillin, 100 μg/ml streptomycin, essential and non-essential amino acids (Gibco), 50 μM β-mercaptoethanol (Sigma), and 10% FBS (Gemini). LAD2 cells were maintained in StemPro-34 serum-free media supplemented with StemPro-34 nutrient supplement (Gibco), 100 ng/ml SCF (Peprotech), 2 mM L-glutamine, 100 U/ml penicillin, and 100 μg/ml streptomycin (LAD2 cell media). THP-1 cells were grown in RPMI 1640 (Gibco) supplemented with 2 mM L-glutamine, 100 U/ml penicillin, 100 μg/ml streptomycin, 20 mM HEPES, 50 μM β-mercaptoethanol (Sigma) and 10% FBS. HUVEC cells were grown in EGM Plus media supplemented with EGM Plus SingleQuots (Lonza).

### Flow cytometry of peripheral blood

Peripheral blood samples from healthy individuals were processed by density gradient centrifugation using Ficoll-Paque Plus (GE Healthcare) to isolate mononuclear cells, which were then cryopreserved. Thawed mononuclear cells were stained with Fixable Viability Dye eFluor 455UV (Thermo Fisher Scientific). We then incubated the cells with MR1 tetramers (NIH Tetramer core) at room temperature for 20 minutes followed by the remainder of surface markers including CD3ε (UCHT1), CD8α (RPA-T8), CD14 (M5E2), CD16 (3G8), CD56 (5.1H11), and Vδ2 (B6) from Biolegend and CD4 (SK3, eBioscience/ThermoFisher), for 20 minutes on ice. Cells were then fixed, permeabilized, and stained for intracellular markers GZMK (GM26E7, Biolegend) and GZMB (GB11, eBioscience/ThermoFisher), using True-Nuclear Transcription Factor Fixation Buffer Set (BioLegend). Data were acquired on a BD Fortessa analyzer using FACSDiva software. Compensation and analysis were performed using FlowJo 10.7.1. Gating of blood cell populations is shown in **Extended Data** Fig. 1a.

### Disaggregation and flow cytometry of synovial tissue

Synovial tissue from arthroplasty or synovectomy samples were cryopreserved in 1 to 5 millimeter fragments for batch processing. Thawed synovial tissues were disaggregated into single-cell suspensions by mincing and digesting with 100 micrograms per milliliter LiberaseTL (Roche) and 100 micrograms per milliliter DNaseI (Roche) in RPMI-1640 (Gibco) for 15 minutes, inverting every 5 minutes^93^. Cells were passed through a 70 micron cell strainer and washed prior to antibody staining using the same protocol described for blood and the same antibodies and clones in addition to antibodies against CD45 (HI30, Biolegend). Gating of synovial cell populations is shown in **Extended Data** Fig. 1c.

### Synovial tissue single-cell RNA-seq analysis

To inspect *GZMK* and *GZMB* gene expression in single-cell transcriptomic data, we used a published synovial tissue CITE-seq dataset from patients with RA and OA^14^ (n = 79). Based on a low-resolution clustering of the data, we subsetted a cluster of T cells (CD3^+^) and a cluster of NK cells and innate lymphoid cells (CD45^+^CD3^-^CD19^-^CD14^-^). Because the original study used CITE-seq, this dataset contained measurements of whole-transcriptome single-cell RNA expression and a panel of 58 surface proteins. We conducted quality control and normalization of the RNA and protein data as described in the original paper. To identify proteins with the most cell-state-specific expression, we coarsely clustered the subsetted data based on RNA expression, and for each protein, measured the Kullback-Leibler (K-L) divergence of cells with high expression of that protein (i.e., normalized expression ≥ 85^th^ percentile) compared to the null distribution of all cells. Then, we scaled proteins with high K-L divergence (n = 25) and highly variable genes to have mean = 0 and variance = 1, and used them for canonical correlation analysis. This strategy projects the cells into a low-dimensional embedding based on both RNA and protein expression. We removed batch effects from this embedding with Harmony^19^ and made a UMAP^94^ with the first 20 canonical variates.

### Integration of multiple scRNA-seq datasets focusing on T cells and innate lymphocytes

To integrate T cells and innate lymphocytes from multiple diseased tissues, we obtained the raw FASTQ files and raw count matrices from the following publicly available scRNA-seq datasets: RA synovial cells from dbGaP^46^ (phs001457.v1.p1) and dbGaP^17^ (phs001529.v1.p1), SLE kidney cells from dbGaP^95^ (phs001457.v1.p1), UC colon cells from Single Cell Portal^15^ (SCP259), CD ileum cells from GEO^16^ (GSE134809), and COVID-19 and healthy bronchoalveolar lavage fluid (BALF) cells from GEO^18^ (GSE145926). For the FASTQs that we obtained, we used Kallisto to map the raw reads to the same kallisto index generated from GRCh38 Ensembl v100 FASTA files. We pseudo-aligned FASTQ files to this reference, corrected barcodes, sorted BUS files, and counted unique molecular identifiers (UMIs) to generate UMI-count matrices. We aggregated all the cells into one matrix and further identified T cells and innate lymphocytes using graph-based clustering and canonical cell lineage gene signatures (**Extended Data** Fig. 2). We performed consistent QC to remove the cells that expressed fewer than 500 genes or with more than 20% of the number of UMIs mapping to the mitochondrial genes.

We applied one unified cross-tissue integrative strategy^96^ to identify functionally meaningful clusters. Specifically, we normalized each cell to 10,000 reads and log-transformed the normalized data. We then selected the top 1,000 most highly variable genes based on dispersion within each donor sample and combined these genes to form a variable gene set. Based on the pooled highly variable genes, we then scaled the aggregated data matrix to have mean 0 and variance 1. We normalized the expression matrix using the L2 norm. To minimize the effect of multiple datasets with different cell numbers during an unbiased scRNA-seq data integration, we performed weighted principal component analysis (PCA) and used the first 20 weighted PCs for follow-up analysis. The summation of the weights for cells from each separate single-cell dataset is equal so that each dataset contributed equally to the analysis. For all cell-type integration, we corrected batch effects on three different levels (sequencing technology, tissue source, and donor sample) simultaneously using Harmony^19^. We used default parameters and also specified theta = 2 for each batch variable, max.iter.cluster = 30, and max.iter.harmony = 20. For Harmony batch correction, we used the same weights from the weighted PCA. We then applied unbiased graph-based clustering on the top 20 batch-corrected PCs. Then, we performed dimensionality reduction using UMAP^94^.

### Immunoblotting

Cell lysates were prepared by lysing cells for 1 h at 4 °C in RIPA buffer supplemented with cOmplete protease inhibitor cocktail (Roche). The cell lysate concentrations were quantified using a BCA protein assay kit (ThermoFisher). Cell-free culture supernatants were precipitated by incubating them with 10% trichloroacetic acid (TCA, Sigma) for 1 h at 4 °C. Cell lysates and precipitated supernatants were separated by SDS-PAGE on 12% or 7.5% Mini-PROTEAN TGX or Criterion TGX gels (Bio-Rad) and transferred onto 0.2 μM PVDF membranes using a Trans-Blot Turbo transfer system (Bio-Rad). Membranes were blocked with 5% nonfat dry milk containing 0.1 % Tween 20 (Bio-Rad) in TBST for 1 h at room temperature, and then incubated with primary antibodies overnight at 4 °C. Membranes were developed using Clarity Western ECL (Bio-Rad) and imaged using a ChemiDoc Touch MP (Bio-Rad).

### Isolation of primary human CD8^+^ T cells for assessment of intracellular and secreted GZMK and GZMB by immunoblot

To isolate human CD8^+^ T cells, peripheral blood mononuclear cells were isolated from healthy donors using Ficoll-Paque Plus density gradient centrifugation followed by magnetic-activated cell sorting using the human CD8^+^ T cell isolation kit (Miltenyi). For assays, CD8^+^ T cells were cultured in RPMI 1640 supplemented with 2 mM L-glutamine, 100 U/ml penicillin, 100 μg/ml streptomycin, 20 mM HEPES, 1 mM sodium pyruvate, essential and non-essential amino acids (Gibco), 50 μM β-mercaptoethanol (Sigma), 5% human AB serum (Gemini) and 30 IU ml^-1^ recombinant human IL-2 (Peprotech). Primary human CD8^+^ T cells were left unstimulated or were stimulated with anti-CD3/CD28 dynabeads (ThermoFisher) at a 1:1 bead-to-cell ratio for 6, 24 and 48 hours. Supernatants were depleted of albumin, immunoglobulins, and other abundant proteins in human serum before precipitation. Cell lysates and precipitated supernatants were immunoblotted against GZMK (EPR24601-164, Abcam, 1:1,000), GZMB (EPR8260, Abcam, 1:1,000), and actin (AC-15, Millipore Sigma, 1:2,000).

### Assessment of GZMK and GZMB expression by stimulated versus unstimulated human CD8^+^ T cells by intracellular flow cytometry

Primary CD8^+^ T cells were purified from cryopreserved human peripheral blood mononuclear cells (N=4) using magnetic bead negative selection (Miltenyi). We cultured 150,000 CD8^+^ T cells in each well of a flat-bottom 96-well tissue culture plate in RPMI with 10% fetal calf serum with or without anti-CD3/CD28 dynabeads at a 1:1 bead-to-cell ratio for up to 4 days. Each day, cells from technical replicate wells were harvested and stained for surface and intracellular markers to measure GZMK and GZMB expression. The flow cytometry panel included Fixable Viability Dye UV455 (eBioscience) and antibodies against CD3 (UCHT1), CD4 (RPA-T4), CD8 (RPA-T8), CD14 (M5E2), GZMK (GM26E7), and GZMB (GB11), all from BioLegend. We also included MR1 tetramers (NIH Tetramer core) to exclude mucosal-associated invariant T (MAIT) cells from analysis. Data were collected on a BD Fortessa flow cytometer and analyzed using FlowJo 10.7.1 software.

### Factor B cleavage assays

To assess CFB cleavage, serum-purified human CFB (Complement Technology, 100 nM) was incubated with CFD (Complement Technology, 100 nM) or increasing concentrations (125 nM, 250 nM, 500 nM, 1000 nM) of recombinant, active human GZMK (Enzo) in the presence or absence of C3b (Complement Technology, 10 nM) in reaction buffer (50 mM Tris-HCl pH 8.0, 150 mM NaCl and 2 mM Mg^2+^ and Ca^2+^) for 4 h at 37 °C. The reaction products were run on SDS-PAGE gels, and cleavage of CFB into Bb was assessed by immunoblot using an anti-CFB antibody (A235, Complement Technology, 1:4,000). The Bb fragment was identified by comparison to the band corresponding to serum-purified Bb (Complement Technology, 15 nM).

### Fluid-phase complement C2, C3, and C4 cleavage assays

To assess C4 cleavage, serum-purified human C4 (Complement Technology, 100 nM) was incubated with increasing concentrations (125 nM, 250 nM, 500 nM) of either recombinant, active human GZMK, recombinant, active human GZMA (Enzo), or serum-purified active human C1s (Complement Technology) in reaction buffer for 4 h at 37 °C. The reaction products were run on SDS-PAGE gels, and cleavage of C4 was assessed by immunoblot using an anti-C4 antibody (22233-1-AP, Proteintech, 1:1,000). The C4b fragment was identified by comparison to the band corresponding to serum-purified C4b (Complement Technology, 30 nM).

To assess C2 cleavage, serum-purified C2 (Complement Technology, 100 nM) was incubated with increasing concentrations (25 nM, 50 nM, 100 nM, 200 nM) of C4 and either GZMK (250 nM) or C1s (250 nM) in reaction buffer for 4 h at 37 °C. The reaction products were run on SDS-PAGE gels and C2 cleavage was assessed by immunoblot using an anti-C2 antibody (A212, Complement Technology, 1:4,000).

To assess whether GZMK can directly cleave C3, serum-purified human C3 (Complement Technology, 100 nM) was incubated with increasing concentrations (125 nM, 250 nM, 500 nM, 1000 nM) of GZMK in reaction buffer for 4 h at 37 °C. As a positive control for generation of C3b, CFB (100 nM) was incubated with C3b (Complement Technology, 10 nM) and CFD (100 nM) along with increasing concentrations (12.5 nM, 25 nM, 50 nM, 100 nM) of properdin (Complement Technology) in reaction buffer for 4 h at 37 °C. The reaction products were run on SDS-PAGE gels and C3 cleavage was assessed by immunoblot with an anti-C3 antibody (A213, Complement Technology, 1:32,000). The C3b band was identified by comparing the cleaved fragments to the band corresponding to serum-purified C3b (15 nM).

To detect C3 convertase formation as assessed by the generation of C3a and C3b, serum-purified C3 (Complement Technology, 100 nM) was incubated with C4 (400 nM) + C2 (400 nM), and either increasing concentrations (125 nM, 250 nM, 500 nM, 1000 nM) of GZMK or GZMA, or 125 nM C1s in reaction buffer for 4 h at 37 °C. The reaction products were run on SDS-PAGE gels and cleavage of C3 into C3a and C3b was assessed by immunoblot using anti-C3 (204869, Millipore Sigma, 1:5,000, for C3b) and anti-C3a (A218, Complement Technology, 1:5,000, for C3a) antibodies. The C3a and C3b bands were identified by comparing the cleaved fragments to the bands corresponding to serum-purified C3a (Complement Technology, 100 nM), C3b (40 nM), and iC3b (Complement Technology, 50 nM).

### Quantitation of complement gene expression in bulk RNA-seq from synovial tissue cells

To quantify local expression of complement genes *C2*, *C3*, *C4A*, and *C4B*, we analyzed a published bulk RNA-seq data from T cells (*n* = 47), B cells (*n* = 29), macrophages (*n* = 46), and fibroblasts (*n* = 45) sorted from disaggregated synovial tissue^46^ from patients with RA or osteoarthritis.

### Assessment of complement C2, C3, and C4 secretion by synovial fibroblasts

To assess for complement C2, C3, and C4 secretion, primary RA synovial fibroblasts were stimulated with 100 ng/ml recombinant IFNG (Peprotech), 1 ng/ml TNF (Peprotech), or both cytokines together for 24 h in fibroblast growth medium containing reduced serum (1% FBS). To detect C2, C3, and C4 secretion by immunoblot, cell culture supernatants were TCA-precipitated and immunoblotted using anti-C2 (E-7, Santa Cruz Biotechnology, 1:50), anti-C3 (204869, Millipore Sigma, 1:5,000), and anti-C4 (JM88-13, Novus Biologicals, 1:1,000 for alpha chain, and A205,Complement Technology, 1:32,000 for beta chain). The C2, C3, and C4 bands were identified by comparison to the bands corresponding to serum-purified C2, C3, C3b, C4 and C4b (50 nM). To detect C2, C3, and C4 secretion by ELISA, cell culture supernatants were analyzed using C2 (ab254501, Abcam), C3 (ab108823, Abcam), and C4 (ab108825, Abcam) ELISA kits.

### Assessment of cleavage of synovial fibroblast-derived complement C3

To detect formation of C3 convertases from fibroblast-derived complement proteins, cell-free supernatants from fibroblasts stimulated for 24 h at 37 °C with a combination of IFNG (100 ng/ml) and TNF (1 ng/ml) in serum-free fibroblast medium were incubated with either GZMK (500 nM), GZMA (500 nM), or C1s (500 nM) for 4 h at 37 °C. The reaction products were run on SDS-PAGE gels and cleavage of C3 into C3a and C3b was assessed by immunoblot using anti-C3 (204869, Millipore Sigma, 1:5,000, for C3b) and anti-C3a (A218, Complement Technology, 1:5,000, for C3a) antibodies. The C3a and C3b bands were identified by comparing the cleaved fragments to the bands corresponding to serum-purified C3a (100 nM) and C3b (40 nM).

### Immunofluorescence microscopy

Formalin-fixed, paraffin-embedded human synovial tissue sections were obtained from the Dana-Farber/Harvard Cancer Center Specialized Histopathology core facility. We removed the paraffin with heat and xylene and then performed antigen retrieval in Tris buffer pH 9 in a steamer for 20 minutes. Slides were blocked with 5% bovine serum albumin (BSA) supplemented with bovine, donkey, and human immunoglobulins (Jackson ImmunoResearch) instead of serum, in order to avoid contamination with external complement components. Slides were then incubated overnight with primary antibodies against GZMK (EPR24601-164, Abcam, 1:500,); C3/C3d (C3D/2891, Neo-Biotechnologies, 1:100, or 7C10, Abcam, 1:10); C5a/C5a des Arg (2942, Abcam 1:20, or 2952, Abcam, 1:50); podoplanin (NZ13, ThermoFisher, 1:150); or CD3 (CD3-12, Abcam, 1:100). Secondary antibodies (Jackson ImmunoResearch) were added at 1:200 dilution. Slides were then treated with Sudan Black B (Sigma) for 15 minutes to diminish autofluorescence. Slides were then stained with Hoechst 33342 nuclear stain (ThermoFisher) and mounted with SlowFade Glass Soft-set Antifade Mountant (ThermoFisher). Images were collected using a Zeiss LSM800 confocal microscope (Confocal Microscopy Core, Brigham and Women’s Hospital) using Zen 2.6 software and analyzed using Fiji (ImageJ2 2.9.0/1.53t).

### LAD2 mast cell degranulation

For degranulation assays, C2 (400 nM) + C3 (100 nM) + C4 (400 nM) were incubated with either GZMK (500 nM) or C1s (500 nM) in LAD2 cell media for 4 h at 37 °C. Then, LAD2 cells were washed and incubated with the reaction products or serum-purified C3 or C3a (100 nM) along with a PE-conjugated anti-LAMP1 antibody (H4A3, Biolegend) for 1 h at 37 °C. Cells were washed twice and further stained with an APC-conjugated anti-CD117 antibody (104D2, Biolegend) to gate on mast cells. After two washes, cells were analyzed on a BD Fortessa FACS analyzer using FACSDiva software for surface LAMP1. Analysis was performed using FlowJo 10.7.1.

### Complement C3b and C4d opsonization assay

Opsonization assays were modified from previously described^97^. Briefly, human umbilical vein endothelial cells (HUVEC, ATCC) were incubated with serum-purified complement components C2 (5nM) + C3 (100nM) + C4 (50nM) with GZMK or GZMA (12.5nM, 25nM, 50nM, or 100 nM) or C1s (100nM) for 4 h at 37 °C in EGM Plus media containing no serum or heparin. In experiments where complete serum was used as a source of complement, HUVEC were incubated with 2% human C1q-depleted serum (Complement Technology) in the presence of GZMK (50 nM, 100 nM, 200 nM, 400 nM) for 2 h at 37 °C. Normal human serum (NHS, Complement Technology) was added to a final concentration of 20% as a positive control. Heparin, when present, was used at a concentration of 0.75 U/mL (Lonza). Cells were analyzed for the presence of surface C3b (3E7/C3b, Biolegend), and/or C4d (12D11, Hycult), and/or GZMK (GM26E7, Biolegend) on a BD Fortessa FACS analyzer using FACSDiva software. Analysis was performed using FlowJo 10.7.1.

### Measurement of surface-bound GZMK and C1s by flow cytometry

To detect binding of GZMK or C1s to the surface of cells, synovial fibroblasts, HUVEC, and THP-1 cells were incubated with GZMK (50 nM) or C1s (50 nM) for 1 h on ice at 4 °C in HBSS with Ca^2+^ and Mg^2+^ (Gibco) containing 10 mM HEPES and 0.5% BSA (Millipore Sigma). Cells were then washed twice and stained with antibodies against GZMK and C1s (M81, Hycult Biotech, 1:10) followed by a PE-conjugated anti-mouse IgG1 secondary antibody (RMG1-1, Biolegend). To detect binding of C1s to the cell surface following addition of the C1 complex, cells were sensitized with mouse IgG2a cell-type specific antibodies (HUVEC and THP-1: CD31, HEC7, Invitrogen, 2 μg; Fibroblasts: HLA-A,B,C, W6/32, Biolegend, 2 μg) or a mouse IgG2a isotype control (MG2a-53, Biolegend) for 15 minute at room temperature. Cells were then washed twice and incubated with purified C1 complex (Complement Technology, 100 nM) or C1s (200 nM) for 30 minutes at 37 °C. Cells were washed twice and stained for C1s followed by a PE-conjugated anti-mouse IgG1 secondary antibody. Cells were analyzed for the presence of surface GZMK or C1s on a BD Fortessa FACS analyzer using FACSDiva software. Analysis was performed using FlowJo 10.7.1.

### Comparison of complement activation by fluid phase vs. membrane phase GZMK

To assess how efficient membrane-bound GZMK is at activating complement, HUVEC cells were incubated with GZMK (200 nM) on ice at 4 °C for 30 minutes to allow GZMK to bind their membrane. Media was then removed, and the cells were incubated with C2 (5 nM) + C3 (100 nM) + C4 (50 nM) for 2 h at 37 °C after which the cell-free supernatants were precipitated with TCA, run on SDS-PAGE gels, and immunoblotted for C2b, C3a, and C4a (the complement activation products that do not associate with surfaces) using anti-C2 (A212, Complement Technology, 1:4,000), anti-C3a (A218, Complement Technology, 1:5,000), and anti-C4a (A206, Complement Technology, 1:1000) antibodies. The C2b, C3a, and C4a bands were identified by comparing the cleaved fragments to the bands corresponding to serum-purified C2b (Complement Technology, 5 nM), C3a (150 nM), and C4a (Complement Technology, 50 nM). For fluid phase reactions, GZMK (200 nM) was incubated with C2 (5 nM) + C3 (100 nM) + C4 (50 nM) for 2 h at 37 °C in the absence of HUVEC cells, and the reactions were precipitated with TCA, run on SDS PAGE and immunoblotted for C2b, C3a, and C4a as above.

### Measurement of C5a generation and bioactivity

To assess generation of C5a, HUVEC cells were cultured with serum-purified C2 (5 nM) + C3 (100 nM) + C4 (200 nM) + C5 (Complement Technology, 50 nM) and increasing concentrations (100 nM, 200 nM, 400 nM) of either GZMK or GZMA in EGM-Plus media containing no serum or heparin for 3 h at 37 °C. Cell culture supernatants were assayed for the presence of C5a by ELISA (BD Biosciences). To assess bioactivity of C5a, Chem-1 cells stably-expressing C5aR1 (Eurofins) were labeled with the cell-permeable calcium binding dye Fluo-5F AM (Invitrogen, 2 μM) in the presence of probenecid (Invitrogen, 2.5 mM). Fluo-5F AM-labeled C5aR1-expressing Chem-1 cells were then incubated with the cell-free supernatants from HUVEC cells cultured as above and the cells were immediately analyzed for calcium flux on a BD Fortessa FACS analyzer using FACSDiva software. Analysis was performed using FlowJo 10.7.1. The average mean fluorescence intensity values obtained from the untreated, Fluo-5F-labeled Chem-1 cells were subtracted from the mean fluorescence intensity values of all other samples.

### Terminal Complement Complex detection

HUVEC cells were incubated with serum-purified C2 (5 nM) + C3 (400 nM) + C4 (200 nM) + C5 (50nM) + C6 (Complement Technology, 50 nM) + C7 (Complement Technology, 50 nM) or C5b,6 (Complement Technology, 8.3nM) in the presence of GZMK or GZMA (25 nM, 100 nM) for 4 h at 37 °C followed by addition of C8 (Complement Technology, 66.7nM) + C9 (Complement Technology, 200nM) for an additional hour. Dynasore (Abcam, 80 μM), when used, was added 4 hours prior to addition of C8 and C9. In experiments where serum was used as a source of complement, HUVEC cells were incubated with 2% human C1q-depleted serum in the presence of GZMK (50 nM, 100 nM, 200 nM, 400 nM) for 2 h at 37 °C. Incubations were performed in EGM Plus media containing no serum or heparin. Cells were evaluated by flow cytometry for surface expression of the TCC using an anti-TCC antibody (aE11, Hycult BioTech, 1:100) followed by an anti-mouse IgG2a-PE secondary antibody (Jackson ImmunoResearch, 1:400) using a BD Fortessa FACS analyzer with FACSDiva software. Analysis was performed using FlowJo 10.7.1.

### Statistics

Statistical analysis was performed using GraphPad Prism 9.5. *P* values were calculated using one-way analysis of variance (ANOVA) with Dunnett’s multiple comparisons tests or a two-way ANOVA with Sidak’s multiple comparisons tests. \**P* < 0.05, \*\**P* < 0.01, \*\*\**P* < 0.001 and \*\*\*\**P* < 0.0001, NS = not significant. Data are mean ± s.d.

## Data availability

CITE-seq single-cell expression matrices and sequencing from the Accelerating Medicines Partnership: RA/SLE Network (Zhang et al^14^) are available on Synapse (https://doi.org/10.7303/syn52297840). The bulk RNA-seq data from sorted synovial cell types, also from the Accelerating Medicines Partnership: RA/SLE Network (Zhang et al^46^), are available at ImmPort (https://www.immport.org/shared/study/SDY998, study accession code SDY998).

---

**Extended Data Fig. 1:**
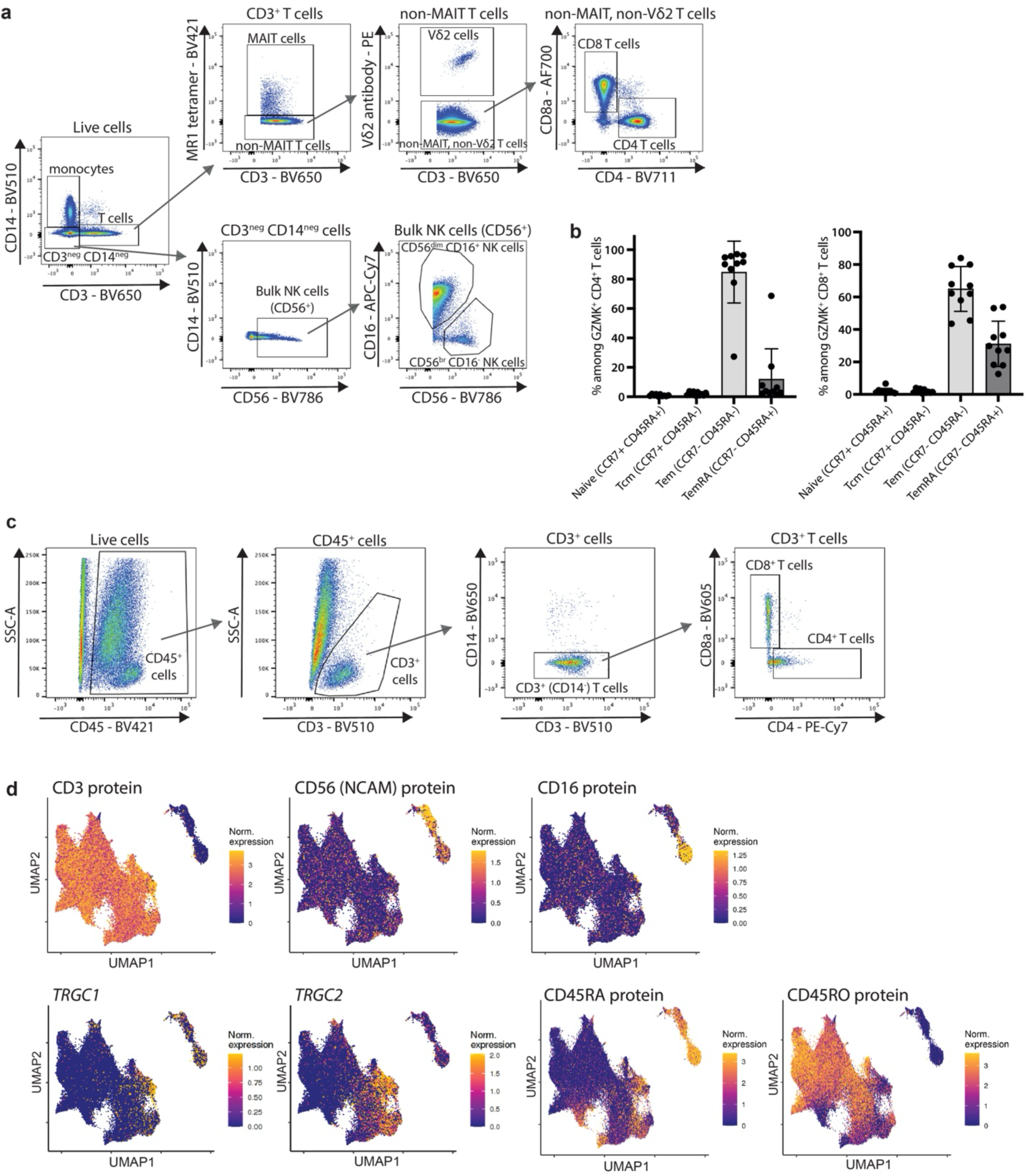
Representative flow cytometry gating and expression of cell-defining markers. **a,** Representative flow cytometry gating of the T cell and NK cell subsets presented in Fig. 1a,b. **b,** Expression of T cell subset markers CCR7 and CD45RA by GZMK^+^ CD4^+^ and CD8^+^ T cells, respectively. Data are mean ± s.d. **c,** Representative flow cytometry gating of CD4^+^ and CD8^+^ T cells in disaggregated RA synovial tissue presented in Figure 1d,e (*n* = 10). **d,** Expression of selected markers in UMAP space for the integrative dataset presented in Fig. 1g.

**Extended Data Fig. 2:**
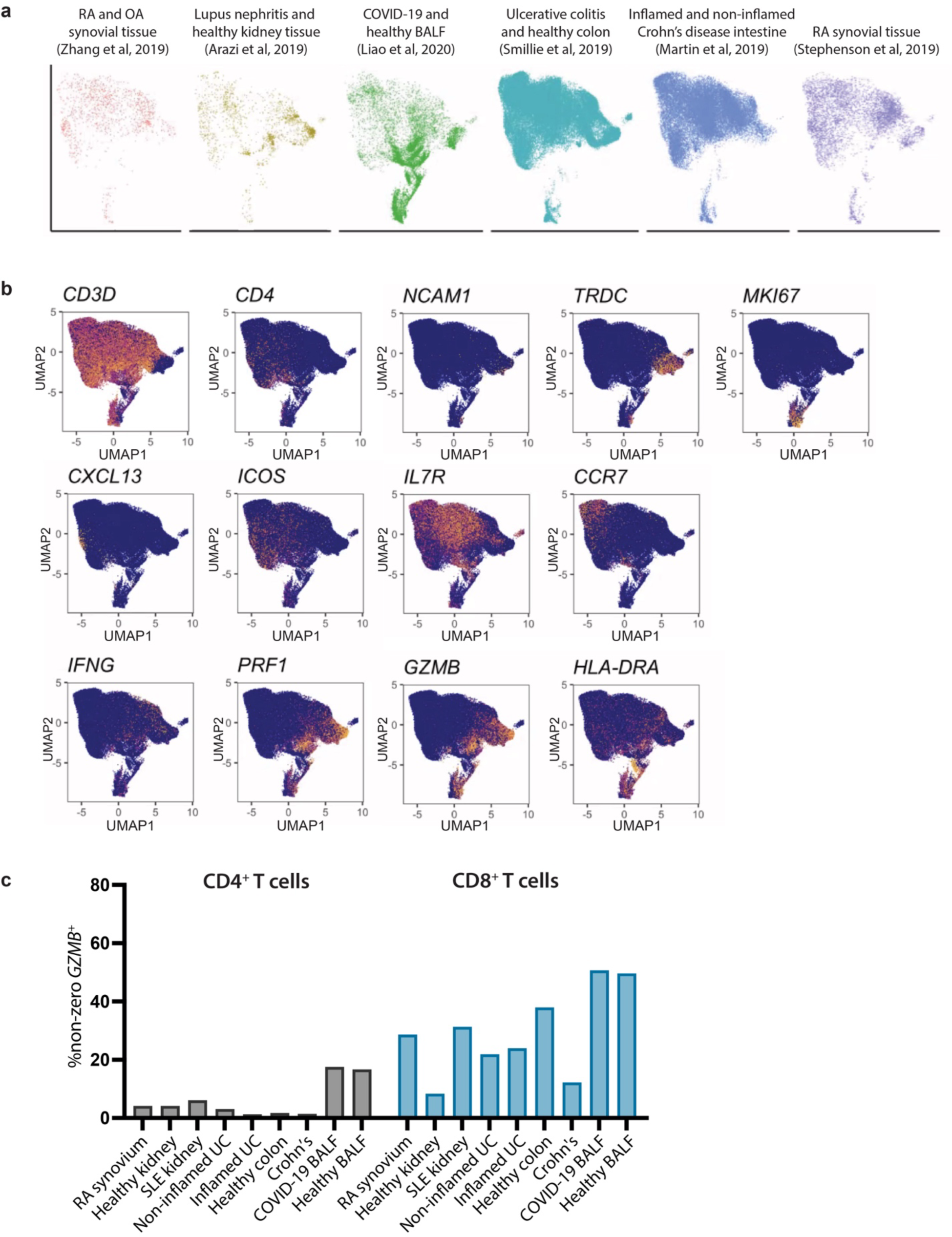
*GZMK* is expressed by CD8^+^ T cells in many different inflamed and non-inflamed tissues. **a,** Contribution of each of the six publicly available single-cell RNA-seq datasets to the integrated dataset of CD4^+^, CD8^+^, and NK cell profiles. **b,** Expression levels of selected genes by cell profiles the integrative dataset in UMAP space. **c,** Percentage of cells in all CD4^+^ T cell clusters (gray columns) or all CD8^+^T cell clusters (blue columns) with detectable *GZMB* gene expression, stratified by tissue and disease source.

**Extended Data Fig. 3:**
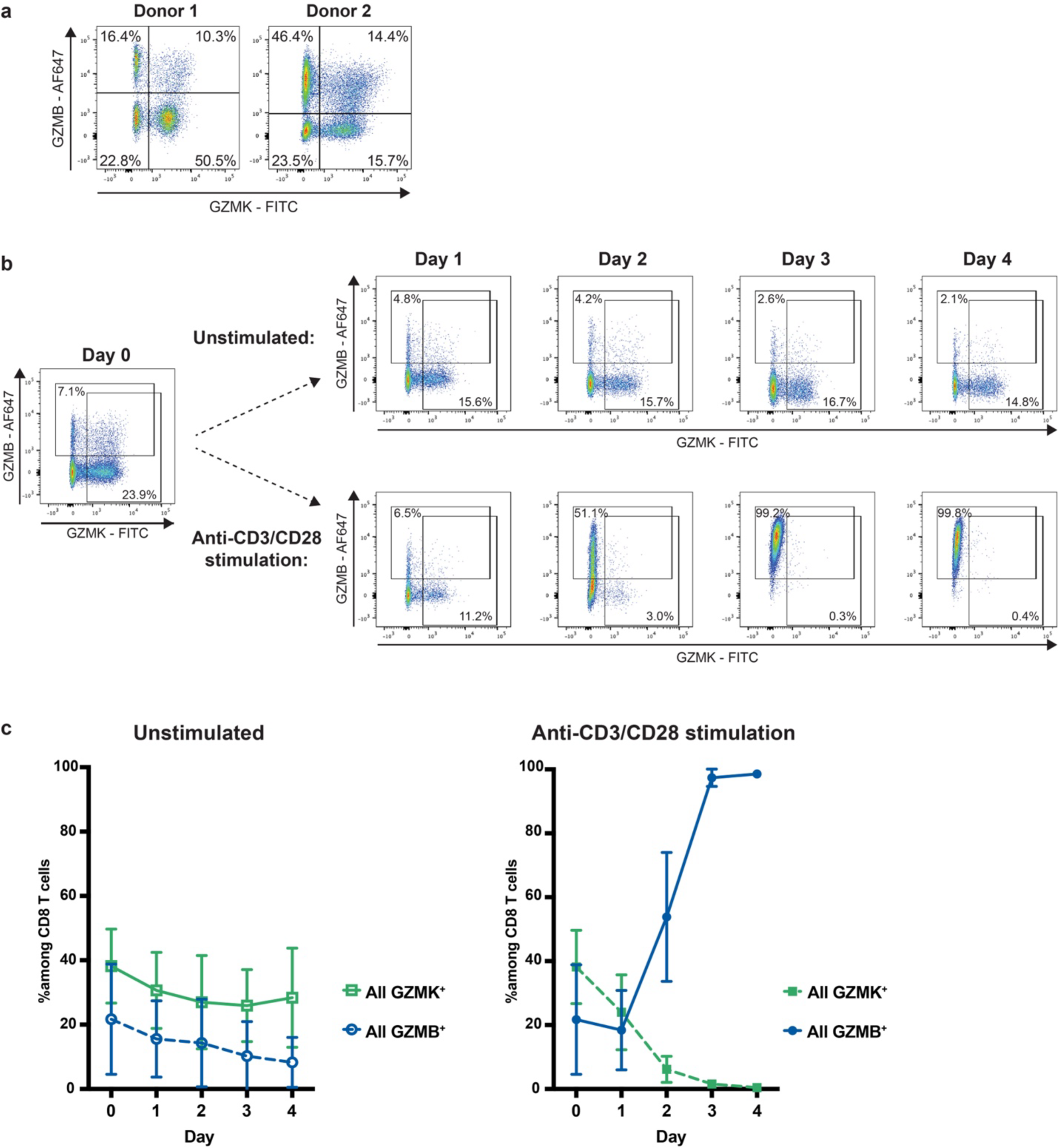
TCR stimulation decreases intracellular GZMK protein expression but increases GZMB in CD8^+^ T cells. **a,** Flow cytometry plots showing intracellular GZMK and GZMB staining among CD8^+^ T cells from the donors used for the immunoblot shown in Fig. 3a. **b,c,** Representative flow cytometry plots and aggregate data showing frequency of intracellular GZMK and GZMB staining among purified primary human CD8^+^ T cells cultured either in media alone (unstimulated) or with anti-CD3/CD28 antibody-coated beads for four days. Data in **c** show mean ± s.d. of four donors from a representative out of four independent experiments.

**Extended Data Fig. 4:**
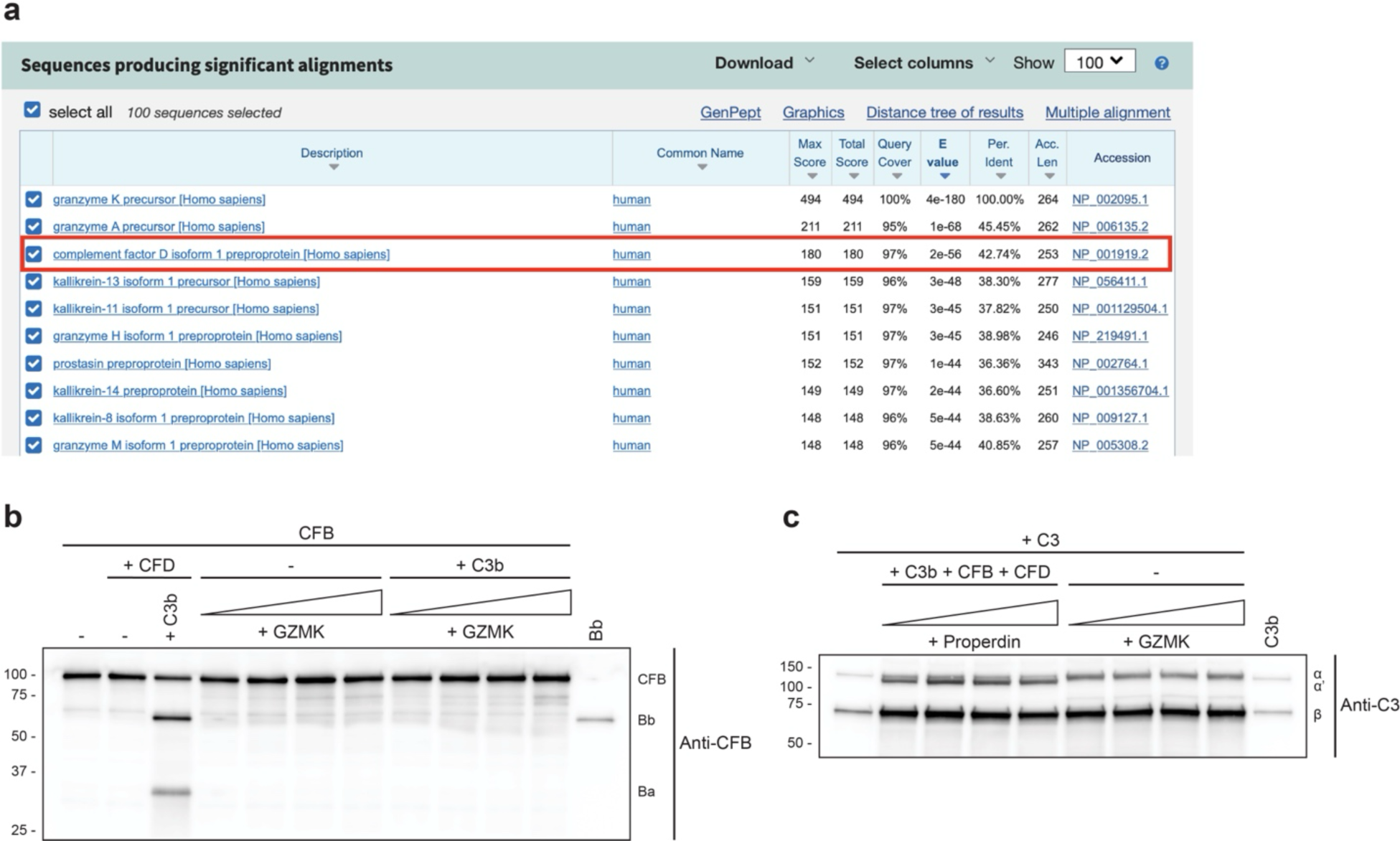
GZMK is highly homologous to complement factor D but does not cleave factor B or C3 directly. **a,** Results of a protein blast comparing the protein sequence of GZMK to all human protein sequences. **b,** Serum-purified complement factor B (CFB) was incubated with either complement factor D (CFD) or increasing concentrations of GZMK in the presence or absence of C3b and cleavage products were analyzed by immunoblot. Serum-purified Bb was used as a control to identify cleavage of CFB into Bb. **c,** Serum-purified C3 was incubated with GZMK and cleavage products were assessed by immunoblot. As a positive control for C3 cleavage, C3 was incubated with C3b + CFB + CFD in the presence of increasing concentrations of properdin. Serum-purified C3b was used to confirm the presence of the C3b cleaved fragment. (**b,c)** Data are representative of three independent experiments.

**Extended Data Fig. 5:**
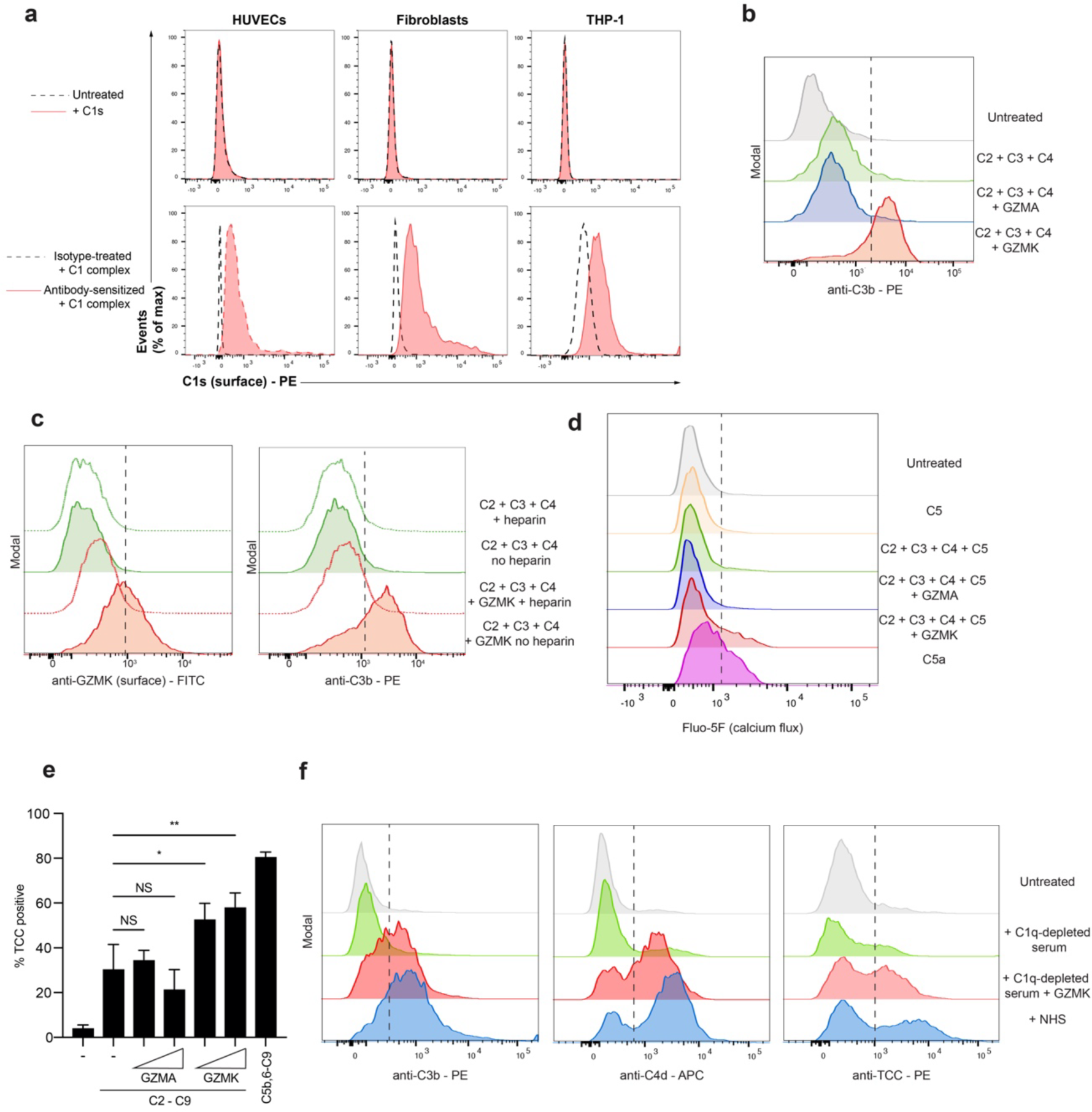
GZMK binds plasma membranes to trigger formation of membrane-bound C3 and C5 convertases. **a**, HUVEC cells, synovial fibroblasts, and THP-1 monocytes were left untreated or were treated with an isotype control or a cell-type specific sensitizing antibody (anti-HLA-A/B/ for fibroblasts, anti-CD31 for HUVEC and THP-1 cells). Cells were then incubated with either C1s or the C1 complex and surface C1s was measured by flow cytometry. **b**, HUVEC cells were incubated for 4 hours with serum-purified C2 + C3 + C4 alone or in combination with GZMK or GZMA, and C3b deposition was measured by flow cytometry. Histograms depict representative data. Aggregate data is shown in Fig. 4c. **c**, HUVEC cells were incubated for 4 hours with serum-purified C2 + C3 + C4 alone or in combination with GZMK in the presence or absence of heparin and surface-bound GZMK and C3b were measured by flow cytometry. Aggregate data is shown in Fig. 4d. **d**, C5aR1-expressing Chem-1 reporter cells labeled with Fluo-5F were incubated with C5, C5a, or the supernatants obtained after incubating HUVEC cells with C2 + C3 + C4 + C5 with or without GZMA or GZMK. Calcium flux was immediately assessed by flow cytometry. Aggregate data is shown in Fig. 4g. **e**, HUVEC cells were incubated with serum-purified C2 + C3 + C4 + C5 + C6 + C7 + C8 + C9 with increasing amounts of GZMK or GZMA in the presence of dynasore to inhibit endocytosis. C5,b-9 was used as the positive control. Terminal complement complex formation (TCC) was then measured by flow cytometry. **f,** Surface deposition of C3b (left), C4d (middle), and TCC formation (right) on HUVEC cells after incubation with C1q-depleted serum and increasing concentrations of GZMK, as measured by flow cytometry. NHS was used as a positive control. Histograms depict representative data. Aggregate data is shown in Fig. 4i. (**a-e**) Data are representative of at least three independent experiments. (**e**) Data are mean ± s.d of three independent experiments. *P* values were calculated using one-way analysis of variance (ANOVA) with Dunnett’s multiple comparisons tests. *P < 0.05, **P < 0.01, NS = not significant.

**Extended Data Fig. 6:**
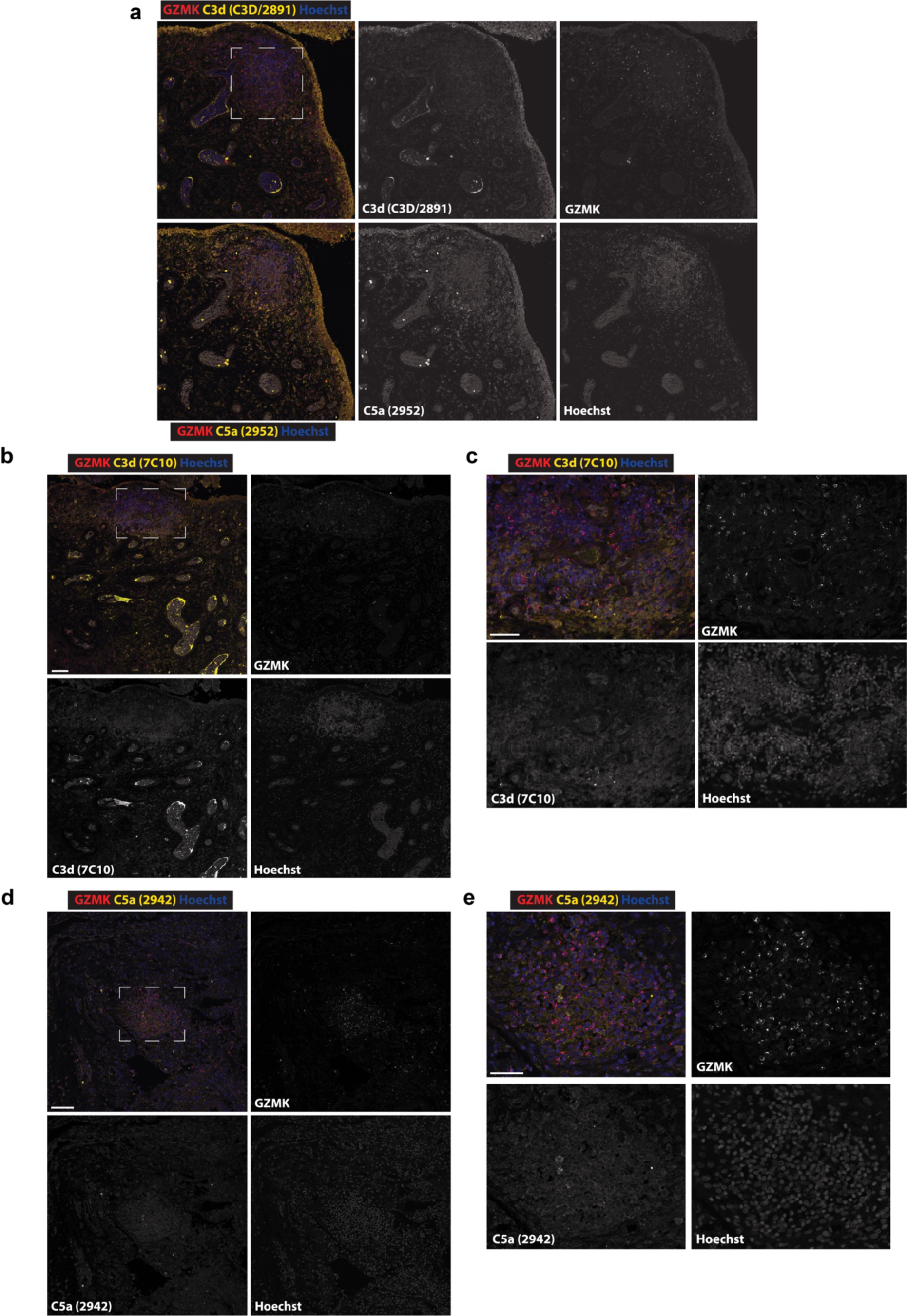
The complement activation products C3d and C5a are abundant in areas rich in GZMK in RA synovial tissue. **a**, Representative image showing immunofluorescence staining of RA synovial tissue with antibodies against C3d (clone C3D/2891, yellow in top composite), C5a (clone 2952, yellow in bottom composite), and GZMK (red) as well as Hoechst nuclear stain (blue). The area inside the dashed box is shown enlarged in Fig. 5. Scale bar is 50 microns. **b**, Representative image showing immunofluorescence staining of RA synovial tissue with antibodies against GZMK (red) and C3d (clone 7C10, yellow) as well as Hoechst nuclear stain (blue). Scale bar is 50 microns. **c**, Enlarged view of area inside box in panel a. Scale bar is 25 microns. **d**, Representative image showing immunofluorescence staining of RA synovial tissue with antibodies against GZMK (red) and C5a (clone 2942, yellow) as well as Hoechst nuclear stain (blue). Scale bar is 50 microns. **e**, Enlarged view of area inside box in panel a. Scale bar is 25 microns. All images in this panel are tiled images collected on a confocal microscope. Data are representative of (**a-c**) three and (**d,e**) two independent experiments.

**Extended Data Fig. 7:**
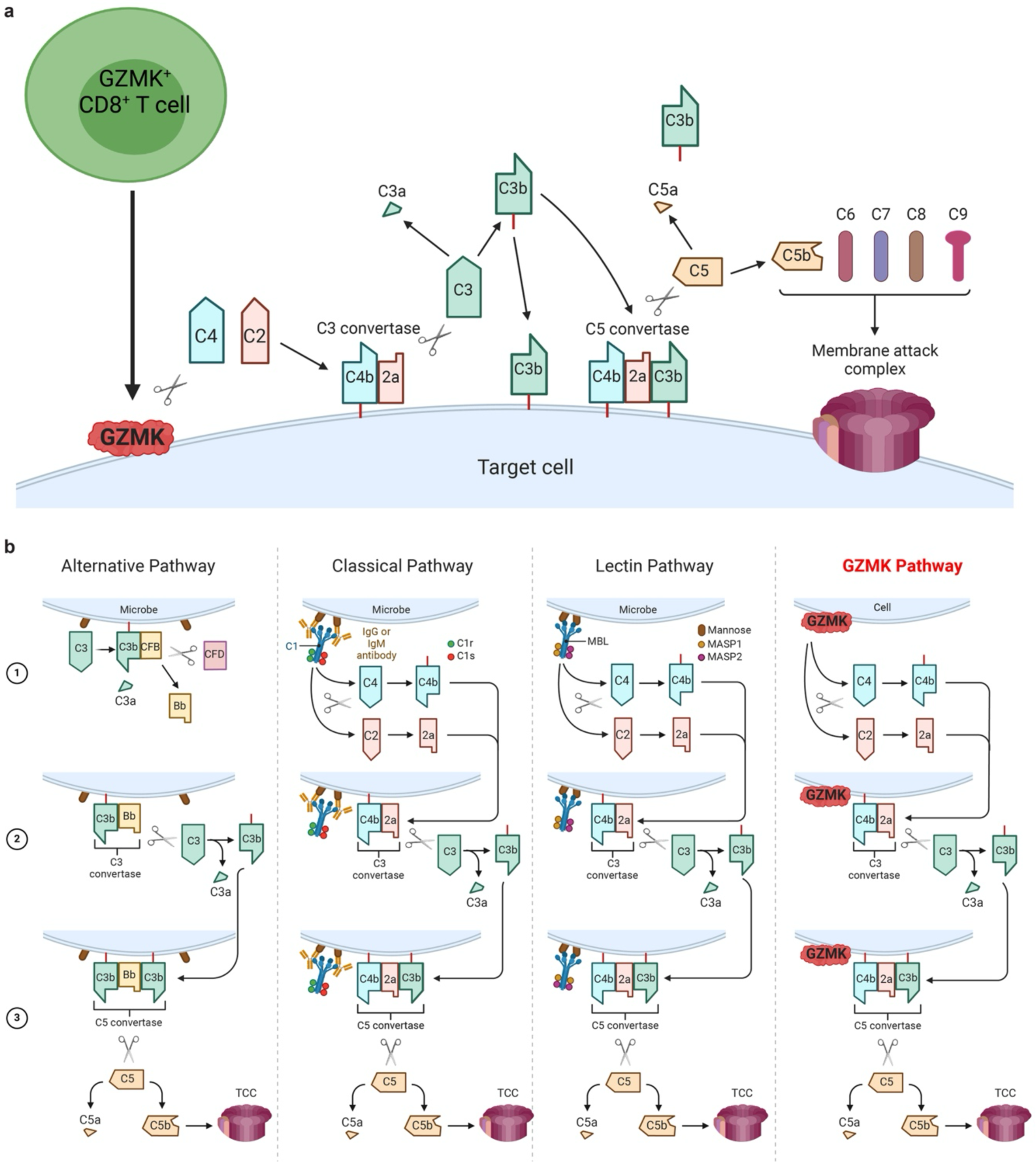
GZMK activates the entire complement cascade. **a.** Model for GZMK-mediated complement activation. *GZMK*-expressing CD8^+^ T cells constitutively release GZMK in the absence of TCR stimulation. Secreted GZMK can bind plasma membranes likely by interacting with heparan sulfate glycosaminoglycans, where it cleaves C4 and C2 generating C4b and C2a. Due to its close proximity to the membrane, newly-cleaved C4b molecules can covalently bind membranes through their exposed thioester, associate with C2a, and form membrane-bound C3 convertases. These C3 convertases can cleave C3 into C3a and C3b. Nascent C3b molecules can opsonize target cells or associate with membrane-bound C3 convertases to form C5 convertases that can cleave C5 into C5a and C5b. C5b molecules associate with C6, C7, C8, and C9 to form a membrane attack complex (MAC) or terminal complement complex (TCC). **b.** Comparison between the alternative, classical, lectin, and GZMK-mediated pathways of complement activation. Complement activation occurs most efficiently on membranes. Soluble pattern recognition receptors like C1q and mannose-binding lectin initiate this process after recognizing danger signals on a surface. This first step results in the activation of initiator proteases like C1s and MASP1, which cleave the complement components C4 and C2. GZMK can direct complement activation to surfaces independently of soluble pattern recognition receptors due to its intrinsic ability to bind negatively charged molecules such as heparan sulfate glycosaminoglycans. Much like C1s, MASP1/2, and CFD, GZMK acts as an initiator protease that cleaves C4 and C2 into C4b and C2a. In the second step, C4b and C2a assemble at the membrane to form active C3 convertases that cleave C3 into C3a and C3b. In the third step, nascent C3b molecules join pre-existing C3 convertases to form C5 convertases that cleave C5 into C5a and C5b, with the latter associating with C6-9 to assemble the TCC/MAC.

